# Mapping and modeling human colorectal carcinoma interactions with the tumor microenvironment

**DOI:** 10.1101/2022.09.13.506996

**Authors:** Ning Li, Qin Zhu, Yuhua Tian, Kyung Jin Ahn, Xin Wang, Zvi Cramer, Ian W. Folkert, Pengfei Yu, Justine Jou, Stephanie Adams-Tzivelekidis, Priyanka Sehgal, Najia N. Mahmoud, Cary B. Aarons, Robert E. Roses, Andrei Thomas-Tikhonenko, Emma E. Furth, Ben Z. Stanger, Anil Rustgi, Malay Haldar, Bryson W. Katona, Kai Tan, Christopher J. Lengner

**Affiliations:** Department of Biomedical Sciences, School of Veterinary Medicine, University of Pennsylvania, Philadelphia, PA 19104, USA; Department of Pediatrics, Perelman School of Medicine, University of Pennsylvania, Philadelphia, PA 19104, USA; Division of Oncology and Center for Childhood Cancer Research, Children’s Hospital of Philadelphia, Philadelphia, PA 19104, USA; Department of Medicine, Perelman School of Medicine, University of Pennsylvania, Philadelphia, PA 19104, USA; Department of Pathology and Laboratory Medicine, Perelman School of Medicine, University of Pennsylvania, Philadelphia, PA 19104, USA; Division of Colorectal Surgery, Department of Surgery, University of Pennsylvania, Philadelphia, PA 19104, USA; Division of Endocrine and Oncologic Surgery, Department of Surgery, University of Pennsylvania, Philadelphia, PA 19104, USA; Division of Cancer Pathobiology, Children’s Hospital of Philadelphia, Philadelphia, PA 19104, USA; Department of Medicine, Vagelos College of Physicians and Surgeons, Herbert Irving Comprehensive Cancer Center, Columbia University, New York City, NY 10032, USA; Abramson Cancer Center, Perelman School of Medicine, University of Pennsylvania, Philadelphia, PA 19104, USA; Institute for Regenerative Medicine, University of Pennsylvania, Philadelphia, PA 19104, USA

## Abstract

The initiation and progression of cancer are inextricably linked to the tumor microenvironment (TME). Understanding the function of specific cancer-TME interactions poses a major challenge due in part to the complexity of the *in vivo* microenvironment. Here we predict cancer-TME interactions from single cell transcriptomic maps of both human colorectal cancers (CRCs) and mouse CRC models, ask how these interactions are altered in established, long-term human tumor organoid (tumoroid) cultures, and functionally recapitulate human myeloid-carcinoma interactions *in vitro*. Tumoroid cultures suppress gene expression programs involved in promoting inflammation and immune cell migration through receptor-ligand interactions, providing a reductive platform for re-establishing carcinoma-immune cell interactions *in vitro*. Introduction of human monocyte-derived macrophages into tumoroid cultures instructs macrophages to acquire pro-tumorigenic gene expression programs similar to those observed *in vivo*. This includes hallmark induction of *SPP1*, encoding Osteopontin, an extracellular CD44 ligand with established oncogenic effects. Taken together, these findings offer a framework for understanding CRC-TME interactions and provide a reductionist tool for modeling specific aspects of these interactions.

## Introduction

Colorectal cancer (CRC) is the third most deadly and fourth most commonly diagnosed cancer globally, with increasing incidence in both developing and developed nations and only minor gains in decreasing mortality rates, primarily among older patients (65+ years)^1,2^. While the reasons underlying the global burden and lack of major therapeutic advances in CRC are complex and multifactorial, a paucity of clinically relevant mouse and human models plays a role. Mouse genetic models have shed light on the molecular events leading to CRC initiation, but are less tractable for modeling aggressive stages of the disease due to the number of driver gene mutations required to establish invasive adenocarcinoma as well as premature mortality resulting from obstruction prior to tumor invasion and metastatic spread. In contrast, human CRC cell lines have been established from late-stage disease, but suffer from years-long culture adaptation, genetic drift, and absence of the complex *in vivo* microenvironment. In the past decade, breakthroughs in our understanding of the intestinal stem cell compartment have enabled, for the first time, the establishment of intestinal epithelial organoid cultures that retain stem cell function and karyotypic normalcy for long periods of time *in vitro*^3^. This knowledge subsequently enabled the establishment of organoids from resected human colorectal cancers^4^.

As with their non-transformed counterparts, colorectal cancer organoids (hereafter referred to as tumoroids) maintain features of tumor tissue architecture observed *in vivo* and, importantly, can predict response to radiation and chemotherapy treatments^5–10^. However, as with traditional 2D cancer cell lines, the lack of tumor microenvironment (TME) components in patient-derived tumor models precludes the development or evaluation of emerging TME-targeted therapies, such as those modulating the activity of cancer-associated fibroblasts or cells of the immune system. Recent advances in air-liquid-interface tumor cultures have begun to address this limitation, enabling modeling of immune checkpoint blockade for example. However, the TME components in tumor air-liquid-interface culture are present only transiently upon culture establishment^11^. Similarly, several studies have sought to understand epithelial-immune crosstalk using organoid co-cultures, including the effects of macrophages on intestinal barrier fidelity, and epithelial-T cell crosstalk ^12,13,14^. Thus, gaining a holistic understanding of carcinoma-TME crosstalk *in vivo* and generating a framework for interrogating specific aspects of this crosstalk is critical for the development and evaluation of therapies aimed at altering this crosstalk.

Recently, several single cell transcriptomic profiles of human CRC have shed light onto the identity of numerous cell populations within the TME^15–17^. Using clustering analysis and trajectory inferences, several tumor-associated macrophage (TAM) populations were identified and postulated to derive from tumor-infiltrating monocyte precursors. These studies hypothesize that, upon association with the TME, macrophages acquire specific states characterized by hallmark expression of genes such as *SPP1* and *C1QC*, and, importantly, do not conform to classical models of M1/M2 macrophage polarization and rather appear to exist as a continuum of states. These studies hypothesized that these TAM states are influenced by numerous factors within the TME, including immune cells, oxygen tension, and nutrient availability. However, insight into the potentially instructive role of carcinoma cells themselves is limited, in part due to the paucity of carcinoma cells in the datasets, as well as the absence of a reductionist experimental model in which the influence of carcinoma cells on macrophage precursors or other cells of the TME can be directly tested.

Here we sought to map putative interactions between human CRC cells and cells of the TME using single cell transcriptomic analyses of treatment-naïve, surgically resected human colorectal adenocarcinomas. In parallel, we established CRC tumoroids and colon organoids, the latter from histologically normal adjacent epithelium. We asked how the selective pressure of *ex vivo* organoid culture alters the carcinoma transcriptome and found a striking suppression of gene expression programs related to carcinoma-TME communication in culture. Using the tumoroid model as a framework for understanding specific carcinoma-TME interactions, we asked whether communication between human macrophages and carcinoma cells could be re-established upon co-culture with tumoroids, given the abundance of macrophages in the TME and their important roles in both tumor suppression and promotion. Remarkably, using this approach we found that interactions with carcinoma cells themselves are sufficient to instruct macrophages to induce pro-tumorigenic TAM identities and gene expression programs, including activation of a hallmark *SPP1+* state. The SPP1+ state acquired by macrophages co-cultured with tumoroids is highly analogous to the SPP1+ state observed in tumor-associated macrophages *in vivo*, relative to macrophages derived from normal adjacent tissue. This SPP1+ macrophage state is associated with tumor immunosuppression and poor prognosis^18,19^, and *SPP1* itself encodes Osteopontin, a well-established pro-oncogenic extracellular matrix (ECM) component and CD44 ligand capable of blunting T cell activation^20^. Taken together, our findings highlight the limitations of carcinoma tumoroid culture models, yet demonstrate that these limitations can be advantageous in reducing a highly complex system in order to functionally interrogate its specific components. Further, we conclude that carcinoma cells themselves are powerful and underappreciated regulators of TAM identity with the ability to induce pro-oncogenic states in human monocyte-derived macrophages.

## Results

### Single cell transcriptomics predicts extensive crosstalk between carcinoma cells and the tumor microenvironment

To begin understanding carcinoma-TME interactions in colorectal cancer, we set out to collect primary tumors, phenotypically normal adjacent colonic tissue, and liver metastases (in rare instances where metastases were surgically resectable concomitant to colon resection). Surgical samples were then processed for single cell transcriptome profiling (scRNAseq) and organoid culture in parallel (**Fig. 1A**). *In toto*, we collected tumors from 17 patients spanning tumor grade, CMS type^21^, and stage (**Fig. 1B-D and Fig. S1**). The majority (15/17) of these tumors were microsatellite stable (MSS), with two exhibiting microsatellite instability (MSI) (**Fig. 1B**). Organoid cultures from tumor and normal adjacent epithelium were maintained for at least 4 passages over a minimum of 2 months, and then subjected to scRNAseq using methodology and reagents analogous to those used for primary tissue (**Fig. 1 C, D** and **Fig. S1**). We captured a total of 38,063 cells from the 15 primary tumors, 11,221 cells from normal adjacent tissues, 5,906 cells from 4 metastatic lesions, and 24,156, and 22,046 cells from *in vitro* cultured normal organoids and tumoroids, respectively. Cell type assignment based on scRNAseq profiles indicated that, as expected, primary samples contained cells of the TME (non-epithelial), while organoid/tumoroid cultures consisted exclusively of epithelial/carcinoma cells (**Fig. 1E**).

**Figure 1.**
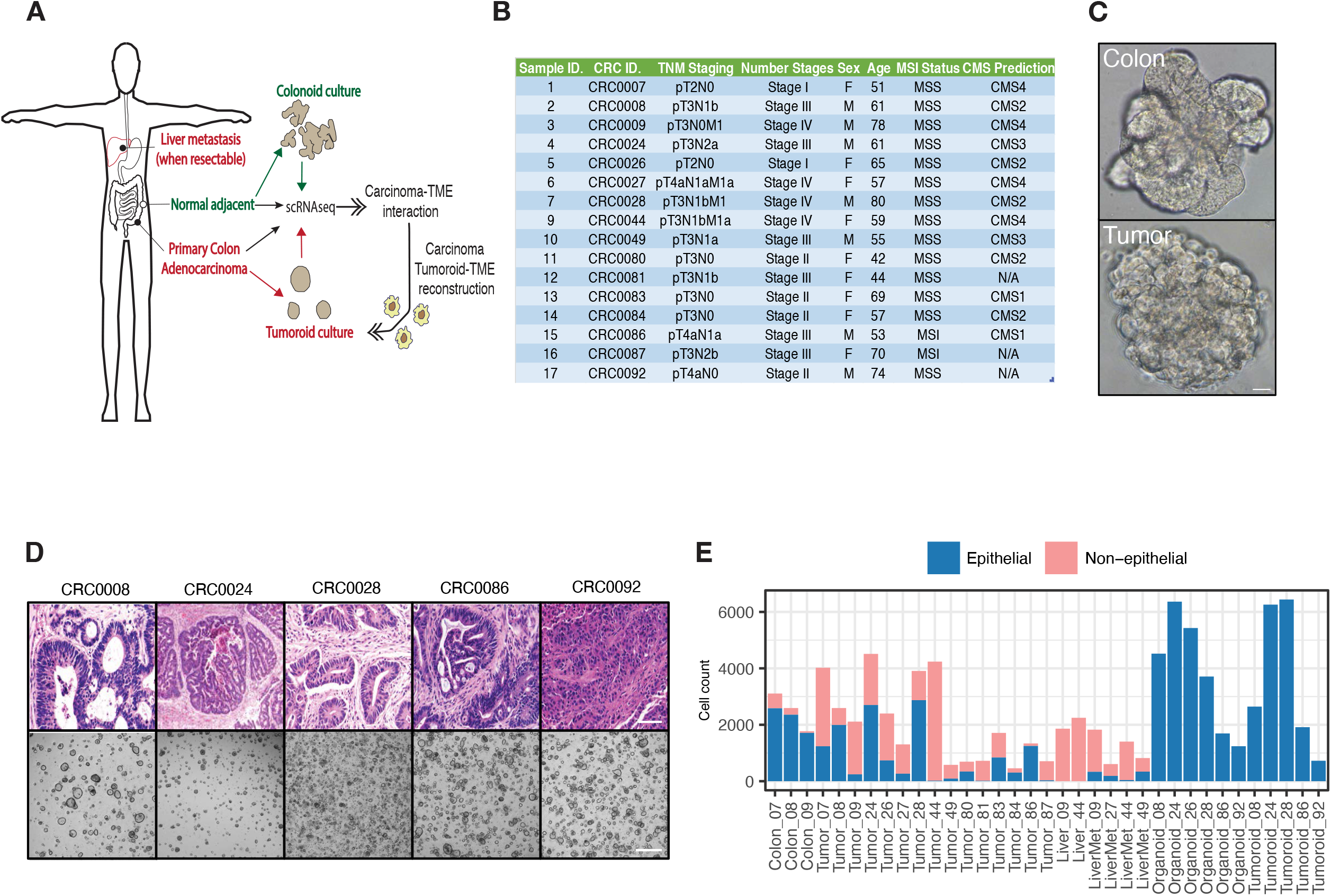
Overview of experimental design: establishment of tumor organoid and single cell transcriptomic datasets. (**A**) Treatment-naïve colorectal adenocarcinomas, liver metastases (when resectable concomitant to primary tumor resection) and normal adjacent colon was subjected to single cell transcriptomic profiling (scRNA-Seq). In parallel, primary tumor and normal adjacent colon samples were also used to seed tumoroid and organoid cultures which were subjected to scRNAseq after culture adaptation. To model TME-carcinoma interactions, healthy human donor-derived monocytes were differentiated into macrophages and co-cultured with organoids or tumoroids and subjected to scRNA-Seq. (**B**). Table providing patient/tumor data. MSI/MSS= Microsatellite instable/stable. TNM staging= Tumor/Node/Metastasis staging. (**C**). Representative brightfield, whole-mount micrograph of normal adjacent-derived colon organoids and primary tumor-derived tumoroids (scale= 50µm). (**D**). Representative hematoxylin/eosin histology for a subset of primary tumors and brightfield micrographs of their cognate 3D tumoroid cultures (scale= 200µm). (**E**) Bar graph showing total sequenced cell count of each sample in scRNAseq datasets, stratified by coarse cell type epithelial (normal or carcinoma) and non-epithelial (normal stroma or TME).

To begin understanding carcinoma-TME interactions in colorectal cancer, we initially evaluated TME composition. Primary tumors contained a variety of cell types, including immune components (macrophages, dendritic cells, T-cells, B-cells, plasma cells, and mast cells), as well as non-immune cell types including endothelial cells, fibroblasts, and myofibroblasts with differing frequencies across patients (**Fig. 2A, B, Fig. S2A**). Immune cells, especially macrophages, T cells, and Plasma/B cells, were present in high abundance in most patients (**Fig. 2B**). In order to understand epithelial-microenvironmental interactions spatially, we also generated cell type annotations in histological sections using CODEX (CO-Detection by indEXing) spatial proteomics (**Fig. 2C, D and S2B-D**). Unsurprisingly, CODEX analyses revealed that in both normal adjacent colon and tumor sections, epithelial (or carcinoma) cells were most likely to make homotypic contacts. However, carcinoma cells had a higher propensity for interaction with cells from the microenvironment relative to their normal counterparts, likely due to a breakdown of normal tissue architecture and tumor infiltration with stromal and immune components (**Fig. 2D**).

**Figure 2.**
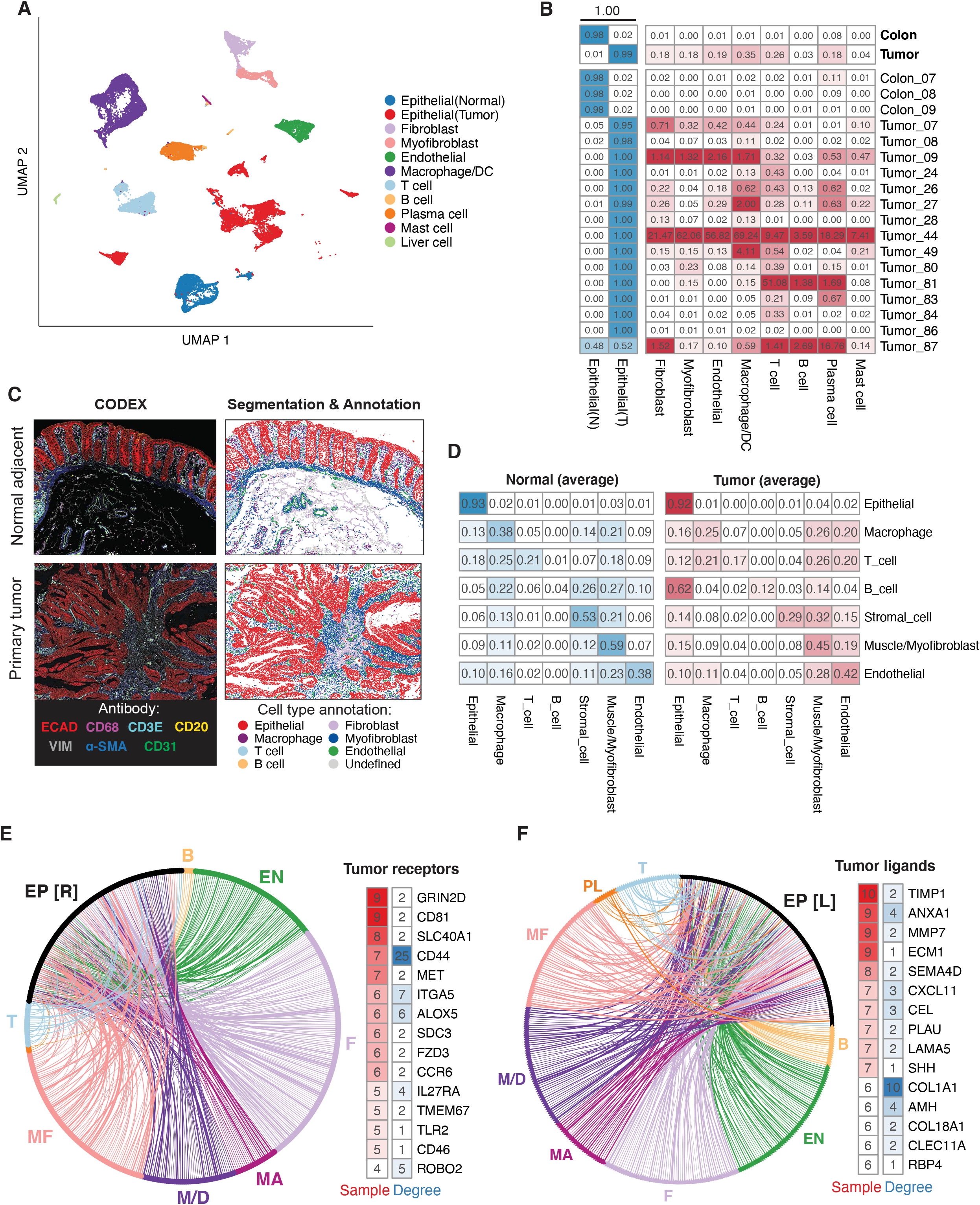
Mapping carcinoma-TME interactions in human colorectal cancer. (**A**) Uniform Manifold Approximation Projection (UMAP) single cell transcriptomes from all cells collected *in vivo*, with cell types indicated. (**B**) Cell type composition in primary tumor and normal adjacent samples. Cell counts of each microenvironment cell type are normalized by total number of epithelial cells. (**C**) Left panel: CODEX image of normal adjacent tissues and primary tumor from patient 86, highlighting seven cell-type markers – ECAD, CD68, CD3E, CD20, VIM, α-SMA and CD31. Right panel: post-segmented image colored by cell type. (**D**) Composition of cell neighborhood based on CODEX image analysis. Each row represents the average cell type composition of a k-nearest-neighbors (k=6) surrounding a particular cell type and the values sum to 1. (**E**) Receptor-ligand interactions up-regulated in primary tumors. Each edge indicates a predicted interaction between a receptor up-regulated in primary tumor carcinoma cells compared to normal adjacent epithelial cells, and a ligand expressed by indicated cell type in the TME. Edge widths indicate the number of patients (samples) in which the receptor is significantly up regulated. Receptors were ranked based on the number of patients with increased expression and the graph degree, which represents the number of ligands from each cell type that communicate with the receptor. EP [R]: receptors expressed on epithelial cells; B: B cells; T: T cells; EN: endothelial cells; F fibroblasts; MF: Myofibroblasts; M/D: Macrophage/dendritic cells; MA: mast cells. (**F**) Same as E, but highlighting the ligands up-regulated in primary tumor carcinoma cells compared to normal adjacent epithelial cells, and corresponding receptors expressed by TME cells. EP [L]: ligands expressed on epithelial cells; PL: plasma B cells.

In order to globally map molecular crosstalk between these diverse cell types, we performed cell-cell communication analyses (see **Methods**), revealing extensive potential receptor-ligand interactions between carcinoma cells and cells within their microenvironment (**Fig. 2E, F, Supplemental Table 1**). Expanding this analysis into metastatic lesions, we also predict numerous carcinoma-TME interactions (**Fig. S3A-C, Supplemental Table 1**). Interestingly, carcinoma cells that metastasize to the liver exhibited similar likelihoods of contact with TME cells as their counterparts in the primary tumor, with some exceptions (**Fig. S3A**). Examining gene expression in metastatic carcinoma cells relative to their primary tumor counterparts, we were able to detect consistent changes in the expression of several genes and pathways in metastases relative to their primary counterparts (**Fig. S3D, E**). This includes the upregulation of pathways governing post-translational control of the canonical Wnt signaling pathway in metastatic carcinoma cells (**Fig. S3E**). Canonical Wnt signaling is the central agonist of colon stem cell self-renewal^22^, and thus this observation may reflect a selection for cancer stem-like cells in metastatic lesions, as has been observed in mouse models of metastatic colorectal cancer^23^.

### Tumoroid culture alters cell type distribution and suppresses gene expression programs associated with epithelial-immune crosstalk

We next sought to understand how removal of carcinoma cells from their *in vivo* tumor environment and introduction into 3D tumoroid culture alters their gene expression programs, particularly as it relates to communication with the TME. We initially focused on understanding the identity of epithelial/carcinoma cells derived from primary tissue versus long-term organoid culture. UMAP-based visualization of single cell transcriptomes and Pearson correlation reveal distinctions not only between normal epithelium and carcinoma cells, but also between primary and cultured cells (**Fig 3A, B**). There was more concordance in transcriptional identity between normal epithelial samples across patients, both in primary and organoid samples, and less concordance amongst tumor/tumoroid samples (**Fig. 3B**). Strikingly, the average transcriptome of cells from tumoroids and organoids are more similar to each other than to their *in vivo* counterparts (**Fig. 3B**). To ask what might account for this, we examined heterogeneity within samples, which revealed a reduction in diversity as carcinoma cells are removed from the primary tumor and maintained in tumoroid culture (**Fig. 3C**). We then examined the fraction of cells with transcriptional identities more similar to crypt base columnar stem cells (SC), transit-amplifying progenitor cells (TA), or mature absorptive colonocytes (CC) across the four sample types (primary colon, organoids, primary tumor, tumoroids) (**Fig. 3D, Fig S4A, B**). As expected, we find that primary normal adjacent colon is enriched in differentiated colonocytes relative to normal organoids, which have greater stem cell and transit-amplifying populations. Interestingly, in cancer, tumoroids also shift towards stem cell and transit-amplifying identity relative to their *in vivo* counterparts (**Fig. 3D**). These observations may be due to the nutrient- and niche cytokine-replete culture conditions driving increased stem cell self-renewal and proliferation relative to the *in vivo* environment.

**Figure 3.**
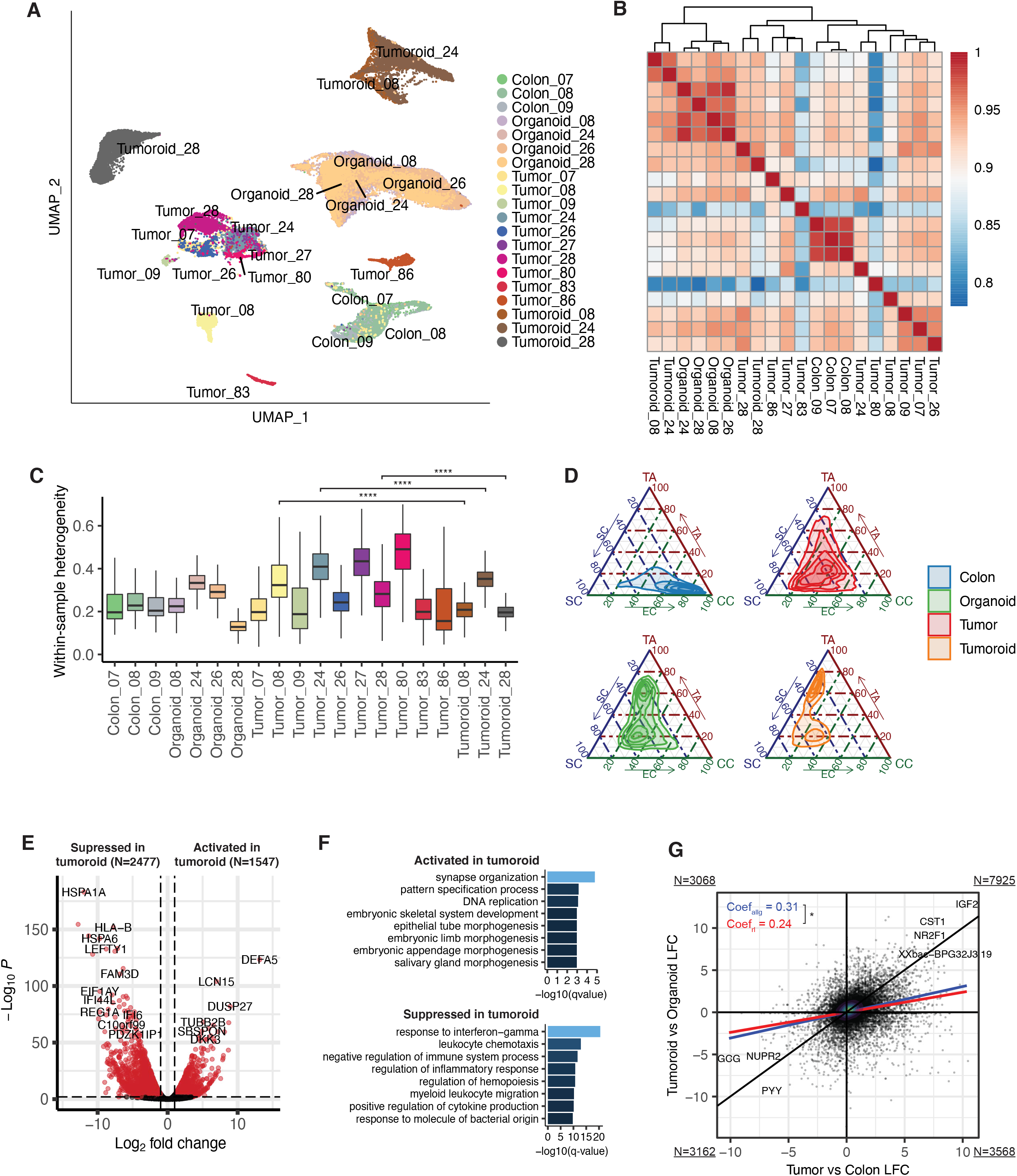
Adaptation to organoid culture suppresses gene expression programs involved in carcinoma-TME communication. (**A**) UMAP of epithelial cells colored by sample. Five tumor samples were excluded from this analysis due to low epithelial cell count. (**B**) Correlation heatmap showing between-sample similarity. Pearson correlations were calculated using average gene expression of epithelial cells for each sample pairs. (**C**) Within-sample heterogeneity across all samples. For each sample, average cosine distance was computed using 500 randomly sampled epithelial cells and displayed as boxplot. (**D**) Ternary plot showing distribution of cells in the cell-type-signature space. Cell type signature genes were derived for each epithelial cell types including stem cell (SC), transit amplifying cell (TA) and colonocytes (CC) using differential expression analysis on normal epithelial cells. Signature scores were calculated using the AUCell package, and linearly rescaled from 0 to 100. 2D kernel density estimation was performed to visualize the cell distribution. (**E**) Volcano plot showing genes activated and suppressed in tumoroid compared to primary tumor based on differential expression analysis. Red point represents genes significantly up- or down-regulated in tumoroid compared to tumor, with FDR ≤ 0.01 and log2 fold change > 1. (**F**) GO functional analysis of DEGs suppressed in tumoroid. -log_10_(q-value) of the top significantly enriched GO terms in biological process were plotted. (**G**) Scatter plot showing correlation of log fold change of each gene *in vivo* (tumor vs adjacent normal, x axis) and *in vitro* (tumoroid vs organoid, y axis). Black line represents y = x (coefficient equal to 1), where the difference between tumor and normal adjacent *in vivo* is the same as their counterparts *in vitro*. Blue line represents fitted linear regression line for all expressed genes, with coefficient equal to 0.31. Red line represents fitted linear regression line for receptors and ligands, with coefficient equal to 0.24. Z test was performed to test the quality of the two regression coefficients^43^ yielding a p-value of 0.024. Number of genes in each quadrant are shown in the corner.

Focusing on carcinoma cells, we next asked how the constraints of the tumoroid culture influence transcriptional identity. We found a greater number of genes suppressed vs. activated in tumoroid culture vs. primary carcinoma cells (**Fig. 3E, Supplemental Table 2**), as well as in organoid culture vs. normal adjacent epithelium (**Fig. S4C, Supplemental Table 2**). Pathway analysis of these downregulated genes indicates that tumoroid/organoid culture primarily suppresses gene expression programs involved in communication with the immune system, particularly those related to leukocyte migration and inflammation (**Fig. 3F, Fig. S4D**). This finding is consistent with the absence of immune cells in these long-term cultures (**Fig. 1D**). Using normal adjacent colon and organoids as control baseline, we compared the log2 fold change of gene expression *in vitro* and *in vivo* and found many genes that are up-regulated in carcinoma cells *in vivo* are also up-regulated *in vitro*. However, the difference between tumoroids and organoids was much smaller *in vitro* than that of their counterparts *in vivo*, suggesting an overall repression of tumor-specific gene expression programs in culture. Interestingly, we found the repression of receptor and ligand expression to be greater than average (p-value = 0.024), indicating a significant impact on gene programs associated with cell-cell communication when moving into organoid culture systems (**Fig. 3G**), Conversely, gene expression programs related to cell division, patterning, and metabolism were generally activated in culture relative to *in vivo* tissue, consistent with the observed shift to stem and progenitor cell states and away from terminally differentiated absorptive states in culture (**Fig. 3D, F, and S4D, E)**.

Given the importance of the TME, and particularly the immune system in colorectal cancer initiation and progression, we wondered how well human tumoroid models represent human primary tumors relative to common *in vivo* models of colorectal cancer in mice with intact immune systems. To this end, we asked how well human tumoroids or various mouse models (including the *Apc*^*min*^ model of familial adenomatous polyposis, the AOM-DSS model of inflammation-driven colorectal cancer, and the APKS model of invasive, metastatic colorectal adenocarcinoma generated by endoscope-guided orthotopic implantation of CRISPR/Cas9-engineered tumoroids with *Apc, p53, Kras*, and *Smad4* mutations into the colonic mucosa of syngeneic mice) recapitulate human primary tumors and tumoroid models. Similar to the human primary tumors, we observed the cancer cell populations shift towards a stem cell-like state in all three mouse models (**Fig. S4 F-H**). However, unlike in human, we did not observe significant population shift towards the proliferative transient-amplifying cell state in the mouse tumors (**Fig. S4H**). Remarkably, human tumoroid transcriptomes are more well-correlated with primary human tumors relative to mouse models, despite the absence of the TME in culture (**Fig. S4 I, J**). Taken together, these findings indicate that the primary effect of long-term tumor organoid culture is a loss of gene expression programs involved in communication with cells of the tumor microenvironment, and that human organoid models may offer advantages in modeling CRC not captured by commonly employed, immune-competent mouse models.

### Human tumors are enriched for pro-tumorigenic macrophage states and depleted of antigen-presenting and pro-inflammatory macrophage states

Recently, several studies have begun elucidating the identity and function of myeloid cells, and particularly macrophages, in the tumor microenvironment of human colorectal cancers^15,16^. Tumor-associated macrophages (TAMs) have been implicated in immune suppression, and macrophage depletion can, in certain tumor types, enhance tumor response to immune checkpoint blockade^24^. TAMs exhibit unique transcriptional gene expression programs that are historically thought to be influenced primarily by the TME (e.g., nutrient availability, hypoxia, fibrotic matrix, and other immune cell types^24,19^). We initially examined the identity and distribution of myeloid populations within our *in vivo*-derived single cell transcriptomic data (**Fig. 4A, B**). We identified a large population of macrophages, including those associated with tumors (primary or liver mets) or with normal tissue (normal colon or liver). We also identified a number of other myeloid dendritic cell (DC) types, including BATF3+ DCs required for effector T cell trafficking and adoptive T cell therapy^25^, CD1c+ DCs which prime cytotoxic T cell responses^26^, Tumor-associated LAMP3+ DCs^27,28^, and LILRA4+ plasmacytoid dendritic cells^29^.

**Figure 4.**
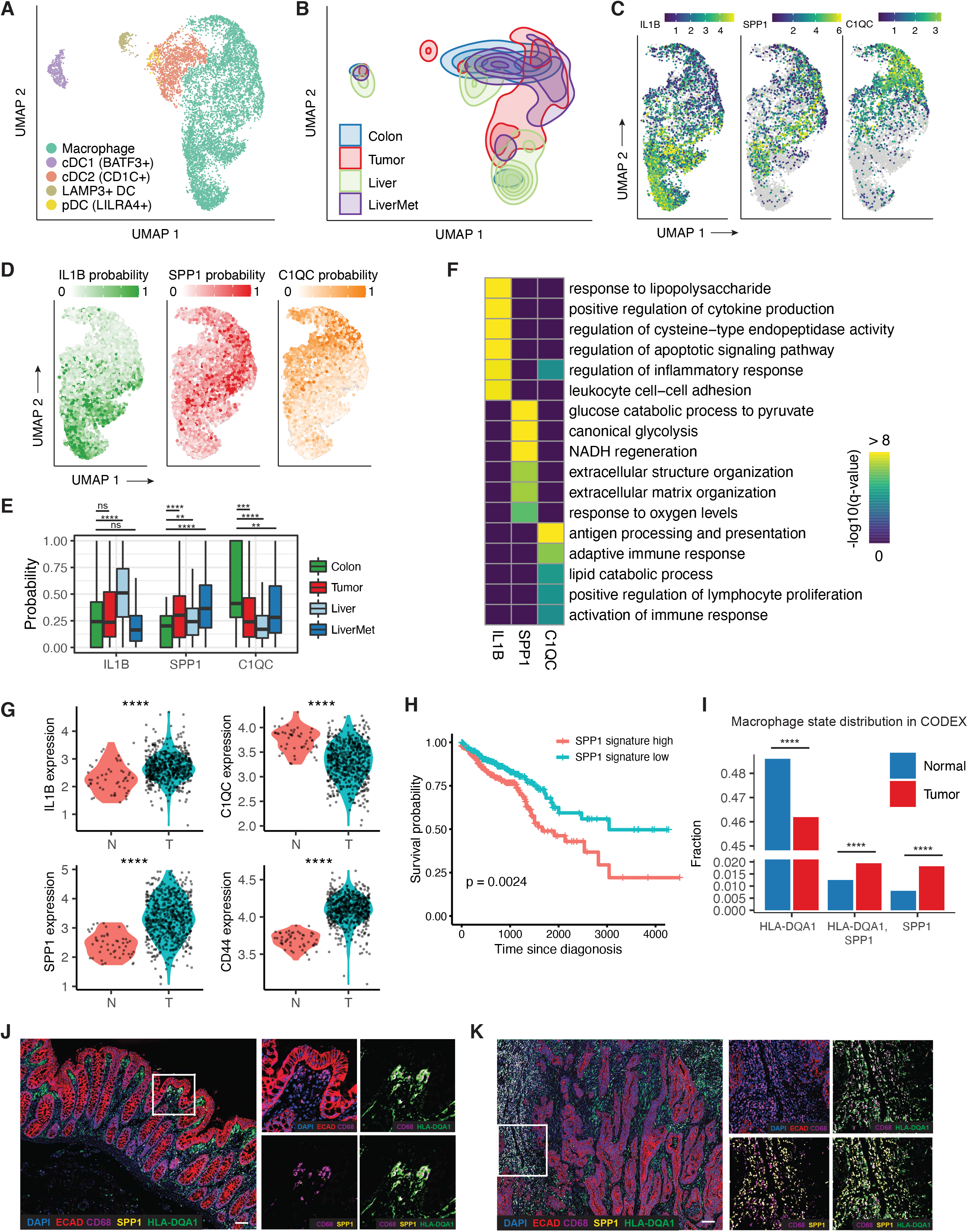
Myeloid cells and macrophage states in human CRC. (**A**) UMAP of myeloid cells present in *in vivo*-derived single cell transcriptomic datasets, colored by indicated cell type. (**B**) UMAP as in (A), but highlighting the distribution of cells across indicated sample types. (**C**) Expression of state-specific hallmark genes in macrophage population. (**D**) Probability of single cells transitioning into each of the macrophage states, calculated by the fateID algorithm. (**E**) Distribution of macrophage state probability across different sample types. Two-sided student’s t test was performed between colon and tumor, colon and liver and colon and liver metastasis. **p≤0.01; ***p≤0.001; ****p≤0.0001; ns, p>0.05. (**F**) Heatmap showing -log_10_(q-value) of top enriched GO terms for each of the macrophage states. (**G**) Expression levels of *IL1B, C1QC, SPP1* and *CD44* in the TCGA colon (COAD) and rectal (READ) datasets. Two-sided student’s t test was performed to compare the expression between tumor and normal samples. ****p≤0.0001. (**H**) Survival analysis on patients stratified by gene signature score of SPP1+ macrophage state. Gene signature score was calculated using the AUCell package and used to group patients into high (≥55th percentile) and low group (≤45 percentile). (**I**) Fraction of HLA-DQA+, SPP1+ and HLA-DQA1+/SPP1+ macrophages in tumor and normal sample in patient 28, 86 and 92 based on CODEX image. HLA-DQA1 antibody was used to label macrophages in the C1QC+ state as these genes were found to be co-expressed based on scRNA-Seq data. Two-sided proportion test was performed to determine if the fraction is significantly different between tumor and normal. ****p≤0.0001. (**J**) Left panel: CODEX image of normal adjacent colon tissue from patient 28, highlighting expression level of ECAD, CD68, SPP1 and HLA-DQA1. Right panels: zoom-in images showing co-staining of markers of macrophage states (scale= 50µm).. (**K**) Analogous CODEX analysis as in (J), but looking at primary tumor tissue from patient 28 (scale= 50µm)..

We next sought to characterize the major macrophage subpopulations present in the data. Using clustering and differential expression analysis (see **Methods**) we identified three major populations: those in an IL-1B+ state, those in an SPP1+ state, and those in a C1QC+ state (**Fig. 4C**). These states are characterized by the co-expression of groups of genes (**Fig. S5A**) and are named for the expression of the hallmark *IL1B, SPP1*, and *C1QC* genes. Examining the distribution of these states in macrophages across sample types revealed that the C1QC+ state is most prevalent in normal adjacent colon tissue where it is associated with antigen presentation and modulation of the adaptive immune response (**Fig. 4D-F, Fig. S5B**). The IL1B+ state is most prevalent in normal liver tissue and is characterized by inflammatory response signatures (**Fig. 4D-F, Fig. S5B**). By contrast, the SPP1+ state is more prevalent in tumors relative to the other states, both in primary tumors relative to normal colon and in liver mets relative to normal liver (**Fig. 4D, E, Fig. S5B**). We predicted extensive receptor-ligand interactions between carcinoma cells and macrophages in each of these states (**Fig. S5C, D**).

The SPP1+ state is characterized by glycolytic gene expression programs, responses to oxygen levels, and gene expression programs related to ECM organization (**Fig. 4F**). Importantly, SPP1+ macrophages have previously been associated with suppression of adaptive immune responses and thus this is considered a pro-tumorigenic state. Further, prior studies suggest that polarization of macrophages toward an SPP1+ state is a result of TME properties, including oxygen tension, the presence of FAP+ cancer-associated fibroblasts, and ECM composition, consistent with our pathway analysis^19,30^ (**Fig. 4F**). Interestingly, *SPP1* itself encodes Osteopontin, a secreted ECM component and ligand for CD44 with known pro-tumorigenic and immunosuppressive activity^18–20,30–32^. Examining expression of *SPP1* and its receptor *CD44* in the TCGA dataset (COAD and READ) reveals concerted upregulation of this receptor-ligand pair in tumors relative to normal tissue (**Fig. 4G**). By contrast, expression of *C1QC* is significantly lower in the TCGA tumors (**Fig. 4G**). Consistent with these reported functions of SPP1 and macrophages in the SPP1+ state, stratification of colorectal cancer TCGA transcriptomes by SPP1 signature enrichment reveals significant reductions in patient survival when tumors have high SPP1 signature (**Fig. 4H**).

Given that these conclusions are drawn largely from single cell transcriptomic analyses, we sought supporting spatial evidence in histological sections of tumor and normal adjacent tissue using co-detection by indexing (CODEX)^33^. We observed macrophages present in histologically normal colonic epithelium were more likely to be HLA-DQA1+ (a proxy marker for the C1QC+ state, **Fig. S5A**), while those present in tumor tissue were more likely to be SPP1+ (**Fig. 4I-K**). Together, these data demonstrate that tumor-associated macrophages are primarily in an SPP1+ immunosuppressive state, and tumors suppress pro-inflammatory and antigen-presenting states.

### Mouse models of colorectal cancer recapitulate macrophage state distributions observed in human tumors

Ultimately, these analyses in human tissue provide valuable insight into the carcinoma-immune TME interactions of colorectal cancer, but platforms for performing functional experiments are severely limited in human systems and mouse models remain the primary platform for performing functional assays targeting carcinoma-TME interactions. We therefore returned to examine the three widely utilized mouse models of colorectal cancer introduced above-the *Apc*^*min/+*^ model of familial adenomatous polyposis, the AOM-DSS model of inflammation-driven colorectal cancer, and the *Apc, p53, Kras, Smad4* quadruple-mutant endoscope-guided orthotopically implanted tumor model of metastatic colorectal cancer (APKS) (**Fig. 5A**). Similar to their human counterparts, these mouse models exhibit similar TME composition with significant myeloid cell infiltration relative to normal mouse colon (**Fig. 5A, B**). Mapping predicted receptor-ligand interactions in the mouse models and cross-referencing these interactions with those observed in humans reveals that the mouse tumors capture only about half of the receptors and ligands expressed in humans (**Fig. 5C, D, Supplemental Table 1**), possibly explaining the limit to DEG recovery in mouse models relative to human tumoroids when compared to primary human tumors seen in Figure 3I. Despite this apparent limitation, however, examining mouse tissues for the presence and distribution of macrophage states identified in human tissue revealed that IL1B+, SPP1+, and C1QC+ states could all be readily identified (**Fig. 5E**). Consistent with human tissue, SPP1+ states were enriched in all three mouse tumor models relative to normal colon, and C1QC+ states were suppressed in all three tumor models (**Fig. 5F**). The pro-inflammatory IL1B+ state was more robustly induced in mouse tumors relative to human, possibly reflecting some fundamental species-specific differences in inflammatory responses, or a limitation of the relatively low number of macrophages captured from normal adjacent human colon samples (**Fig. 5F**). Thus, while these mouse tumor models do not perfectly reflect the cell type distributions and putative carcinoma-TME communication we observe in human, they appear to largely recapitulate macrophage state changes in response to malignancy.

**Figure 5.**
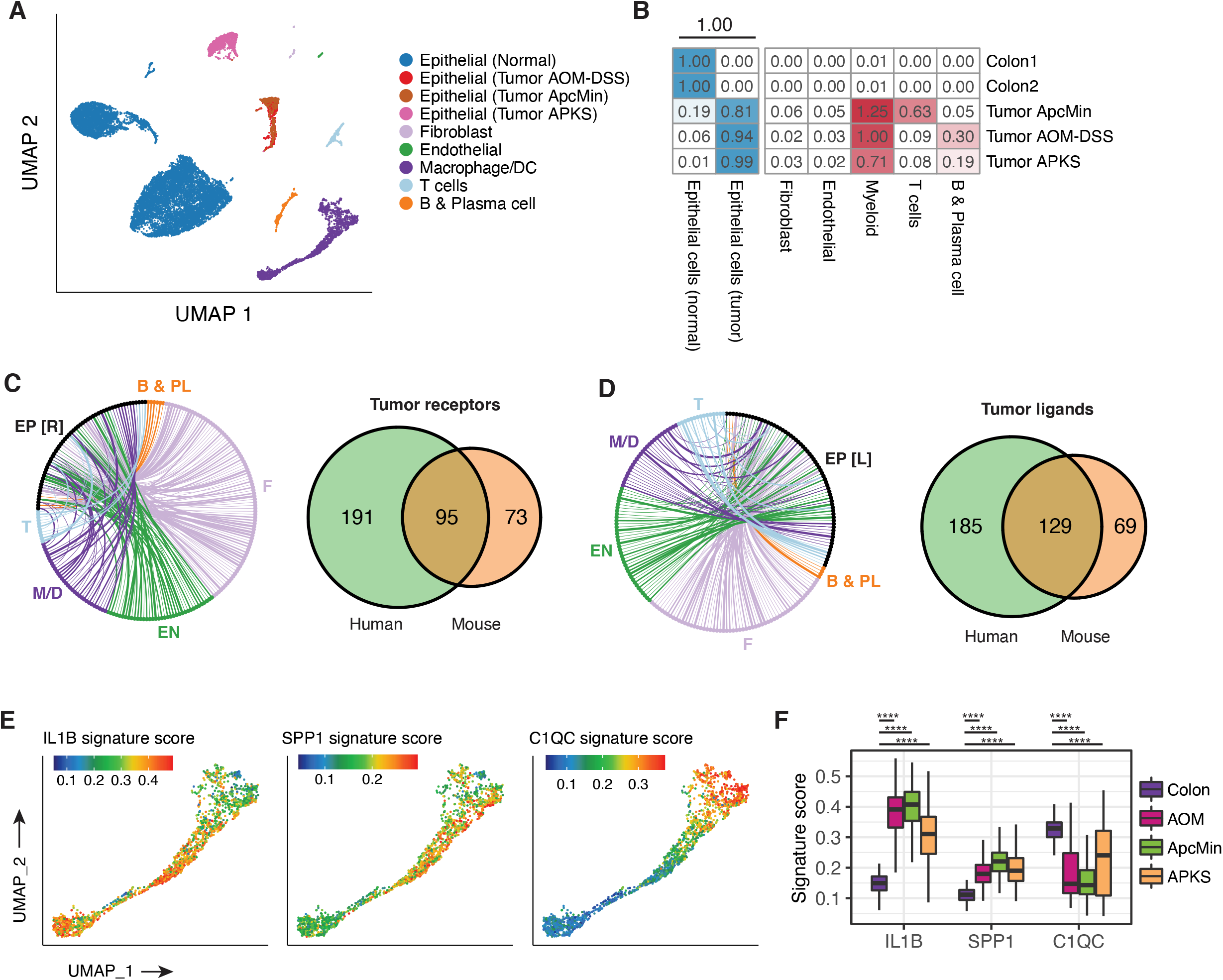
Mouse colorectal cancer models recapitulate human tumor-associate macrophage states. (**A**) UMAP of all cells from mouse CRC models, colored by cell types. (**B**) Cell type composition in mouse CRC models (AOM-DSS inflammation driven colorectal cancer, Apc^min/+^ model of familial adenomatous polyposis, and APKS model employing endoscope-guided orthotopic implantation of engineered tumoroids into the colonic mucosa of immune-competent mice) and normal control colon. Cell counts for each microenvironmental cell type are normalized by total number of epithelial cells. (**C**) Receptor-ligand interactions up-regulated in mouse CRC models. Each edge indicates communication between a receptor up-regulated in mouse tumor epithelial cells vs. normal colon epithelial cells, and a ligand expressed by TME cells. Edge widths indicate the number of CRC models in which the gene is differentially expressed. Venn diagram on the right indicate the overlap of tumor-specific receptors between mouse and human. EP [R]: receptors expressed on epithelial cells; B & PL: B and plasma B cells; T: T cells; EN: endothelial cells; M/D: Macrophage/dendritic cells. (**D**) Same as (C), but highlighting the ligands up-regulated in mouse tumor epithelial cells compared to normal colon epithelial cells, and corresponding receptors expressed by TME cells. EP [L]: ligands expressed on epithelial cells. (**E**) UMAP of macrophages colored by a score that measures the extent to which each single-cell transcriptome is enriched for genes specific to each of the macrophage states observed in human. Signature genes were derived by differential expression and were converted to mouse orthologs. Gene set enrichment scores were computed using the AUCell package^44^. (**F**) Distribution of signature scores for macrophage states in normal mouse colon and mouse CRC models.

### Carcinoma cells instruct macrophages to enter pro-tumorigenic immunosuppressive states

The TME has been extensively implicated in dictating macrophage polarization within tumors, however the degree to which carcinoma cells themselves directly influence macrophage identity remains largely unexplored, particularly in colorectal cancer. We therefore set out to address the degree to which carcinoma cells influence macrophage polarization by taking advantage of the tumoroid model system, free of TME components and absent of the gene expression programs underlying TME-carcinoma crosstalk. To this end, we generated human monocyte-derived macrophages via stimulation with human M-CSF, then introduced these macrophages into either normal organoid or tumoroid cultures, followed by single cell transcriptomic profiling of both the macrophage and epithelial components of the cultures (**Fig. 6A**). In response to introduction of macrophages into tumoroid cultures derived from patient 28 (Stage IV), we observed a clear shift in macrophage polarization toward the SPP1+ state (**Fig. 6B, C**). This shift was consistently seen in macrophages co-cultured with tumoroids derived from three different patients (**Fig. 6C**). There was also modest induction of the IL1B+ state, concomitant to a suppression of the C1QC+ state, largely reflecting the *in vivo* states observed in tumor tissue vs. normal adjacent observed in Figure 4 (**Fig. 6C**). Interestingly carcinoma cells from patient 28 did not reciprocally respond to the presence of macrophages, suggesting that perhaps extended culture may result in epigenetic repression of transcriptional programs responding to the presence of TAMs, and/or the low levels of the Osteopontin receptor CD44 in this tumoroid line (**Fig. 6D)**. By contrast, tumoroid cultures from patient 86 (Stage III) and 92 (Stage II) were responsive to macrophage presence and expressed higher levels of CD44 relative to the non-responsive carcinoma cells from patient 28 (**Fig. 6C-E**).

**Figure 6.**
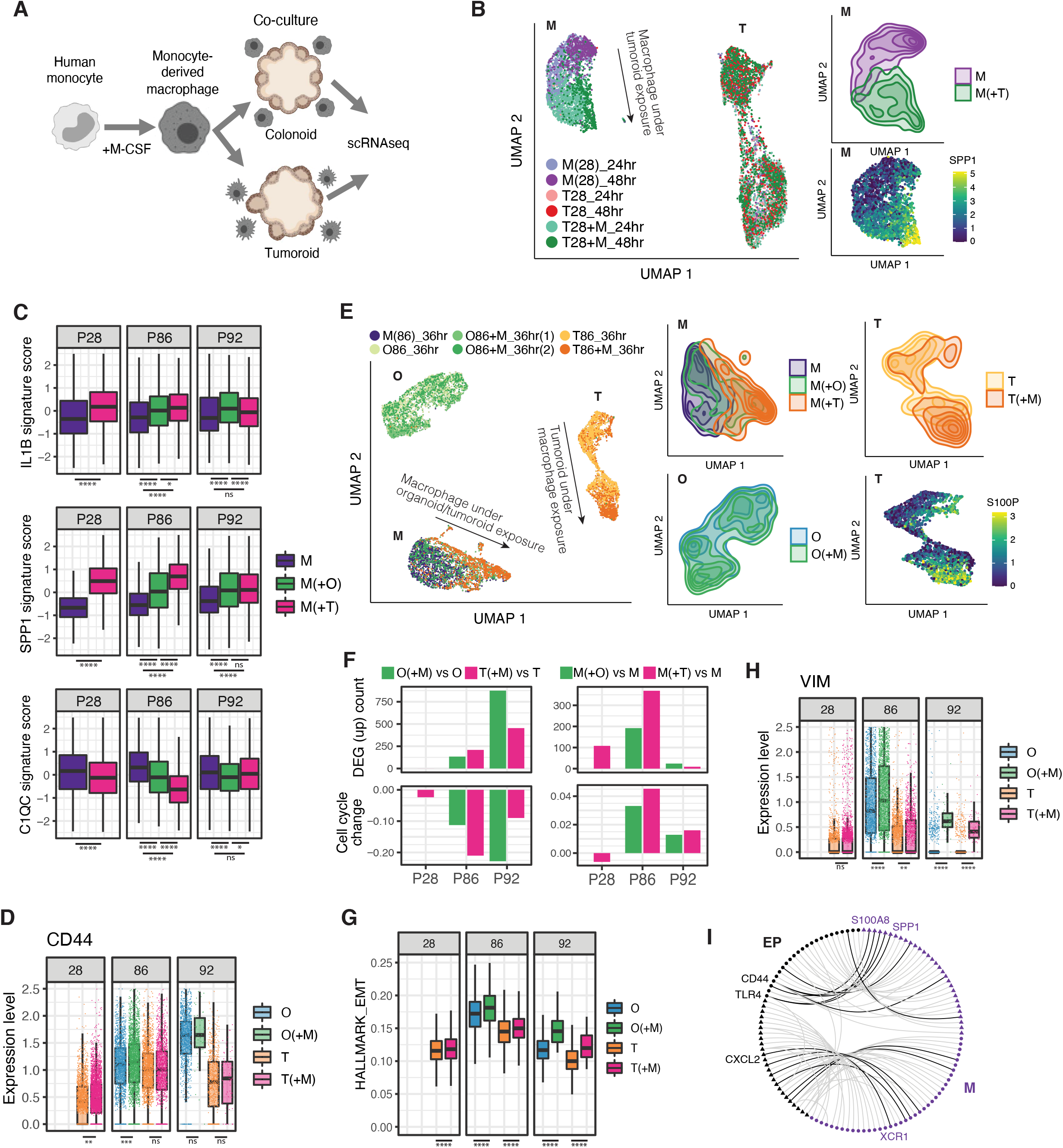
Reconstructing carcinoma-macrophage interactions in tumoroid culture. (**A**) Illustration of tumoroid/organoid-macrophage co-culture assay. (**B**) UMAP of the macrophage and tumoroid cells from Patient 28 before and after co-culture. The top right panel highlights the distribution of macrophage before and after co-culture. The bottom right panel shows the expression of SPP1 in macrophages. M: macrophages; T: tumoroid cells. (**C**) Distribution of signature score for macrophage states before and after exposure to organoids or tumoroids. Data are collected from three independent co-culture experiments with cells from different patients. One-way ANOVA and Tukey’s ‘Honest Significant Difference’ test was used to compare the average signature scores. *p≤0.05; ****p≤0.0001; ns: p>0.05. (**D**) Boxplot showing the expression level of CD44 across condition and patients. One-sided t test was performed with the alternative hypothesis that CD44 expression increased upon co-culturing with macrophage. ***p≤0.001. **p≤0.01; ns: p>0.05. (**E**) UMAP of the macrophages, tumoroid and organoid cells from Patient 86 before and after co-culture. First three panels on the right highlight the distribution of cell states of macrophages (M), tumoroids (T) and organoids (O) before and after co- culture. Last panel shows expression of S100P, a gene up-regulated in tumoroid cells upon co-culture. (**F**) Top left panel shows number of significantly up-regulated genes in each organoid and tumoroid in response to the macrophages. Bottom left panel shows the change in the fraction of organoid/tumoroid cells entering S, G2 or M phase upon co-culture. Right panels show the same set of statistics, but for macrophages before and after co-culture. (**G**) EMT signature score of organoids and tumoroids with or without macrophage co-culture. One-sided t test was performed with the alternative hypothesis that EMT score becomes higher upon co-culturing with macrophage. ****p≤0.0001. (**H**) Expression level of VIM in organoids and tumoroids with or without macrophage co-culture. One-sided t test was performed with the alternative hypothesis that VIM expression increased upon co-culturing with macrophage. ****p≤0.0001. **p≤0.01; ns: p>0.05. (**I**) Recapitulating *in vivo* receptor-ligand interactions in organoid co-cultures. The edges in the graph indicate receptor-ligand interaction between macrophage and tumor epithelial cells *in vivo*. Edges colored in black indicate the R/L interaction up regulated upon co-culture. Circles indicate receptors and triangles indicate ligands.

The clearest consequence of exposing macrophages to carcinoma cells in tumoroid culture was polarization toward an SPP1+ state. The *SPP1* gene product Osteopontin is an extracellular matrix protein with pleiotropic signaling functions. While Osteopontin produced by TAMs contributes to blunting the adaptive immune response to cancer, it has a wide range of cancer cell-autonomous functions including the promotion of an epithelial-to-mesenchymal transition (EMT) linked to increased cancer stem cell properties and metastatic proclivity downstream of its interactions with CD44 and integrin αvβ3^34,32^. We therefore examined the nature of the response of carcinoma cells to the presence of macrophages in co-culture. Macrophage introduction resulted in varying degrees of differential gene expression in patient-derived tumoroids and normal adjacent organoids, and, interestingly, was generally associated with decreased cell cycling of carcinoma cells, but increased cycling of macrophages (**Fig. 6F**). This was concomitant to an increase in EMT signature gene expression in tumoroids and organoids upon co-culture with macrophages (**Fig. 6G**), with clear up-regulation of the mesenchymal marker Vimentin in patient 86 and 92 (**Fig. 6H**). This finding is consistent with a model whereby carcinoma-induced macrophage SPP1 promotes EMT, potentially providing a mechanistic basis for the previously observed link between SPP1 expression and colorectal cancer metastasis^16,34^

Ultimately, we asked to what extent introduction of macrophages into tumoroid cultures recapitulated the macrophage-carcinoma receptor-ligand crosstalk observed *in vivo*. Superimposing *in vitro* receptor-ligand interactions upon the full interactome between carcinoma cells and macrophages *in vivo* (**Fig. 6I**, light lines) reveals that a subset of these interactions (**Fig. 6I**, dark lines) are re-established in culture, including the SPP1-CD44 interaction, among others. Taken together, this study demonstrates that human tumor-derived organoid cultures present limitations for understanding crosstalk between carcinoma cells and the TME, as many of these transcriptional programs are silenced *in vitro*. However, this limitation also offers an opportunity for focused reconstruction of specific carcinoma-TME interactions, enabling a reductionist approach to the functional evaluation of these interactions in what is otherwise a highly complex system *in vivo*.

## Discussion

The recent co-emergence of organoid culture systems and single cell transcriptomics offers unprecedented opportunities for understanding the complex communication between cancer cells and the cells within their microenvironment. Here, we map putative carcinoma-TME interactions from scRNAseq data derived from treatment-naïve colorectal adenocarcinoma, ask how the gene expression programs underlying these interactions are altered in established tumor organoid cultures, and address whether these interactions can be re-established and functionally tested through introduction of specific TME components (macrophages) into the organoid cultures.

The advent of organoid culture, and subsequently tumor organoid (tumoroid) culture, holds promise for the advancement of patient-specific personalized medicine^35^. Recent studies indicate that patient-derived tumoroids can successfully predict response to radio- and chemotherapies^5–10^, and tumoroids have been employed to study the development of tumor-reactive T cells^36^ or the response to immune checkpoint inhibitors in coculture systems^11^. In the context of colorectal cancer, air-liquid interface culture systems support survival of cells of the TME in conditions that favor carcinoma cell growth for several weeks, ultimately non-carcinoma cells do not self-renew and are lost over time^11^.

We initially set out to ask how colorectal carcinoma cells and normal colonic epithelium respond to adaptation to the three-dimensional organoid culture. Our findings were not surprising: organoid culture favors a population shift towards proliferative stem and progenitors, likely resulting from the nutrient and growth-factor replete medium and non-hypoxic environment (our study is conducted using roughly physiological oxygen concentrations of 5%). Long-term organoid culture also led to suppression of gene expression programs involved in cell-cell communication (receptor-ligand pairs) and in immune cell migration and inflammation, a reflection of the absence of TME in these cultures. While these features of organoid culture may prove limiting for some studies (e.g., immune therapies targeting a cell-surface receptor that may become suppressed in tumoroid culture), this reductionist system enables interrogation of the functional consequences of individual TME-carcinoma interactions.

To this end, we focused on the macrophage population as it is a major component of the TME and has been associated with both pro- and anti-tumorigenic activities. Our profiling of macrophages in treatment-naïve CRC and normal adjacent colon was consistent with that of two recent studies employing single cell transcriptomic analyses of colorectal cancers^15,16^. We observed three predominant macrophage states: an antigen-presenting C1QC+ state predominant in the normal colon, a pro-inflammatory IL1B+ state predominantly found in the normal liver, but also in normal colon and adenocarcinoma, and finally, a SPP1+ state enriched in tumors. Reassuringly, these same states were identified in three mouse models of CRC, with roughly similar distributions to the human tumors, highlighting the value of immune-competent mouse models of human cancer.

*SPP1* encodes the extracellular matrix component Osteopontin, which acts as a CD44 ligand and has pleiotropic roles in tumor biology, most notably suppression of T cell activation^9^, and the SPP1+ macrophage state is associated with immune evasion and metastatic proclivity^18,19,37^. Prior studies suggest that polarization of macrophages toward this SPP1+ state is a result of TME properties, including oxygen tension, the presence of FAP+ cancer-associated fibroblasts, and ECM composition, consistent with our pathway analysis of SPP1+ macrophages^19,30,33^. Recent single cell transcriptomic studies hypothesize a circulating monocyte origin for these SPP1+ TAMs^33^. Using the TME-free tumoroid cultures, we were able to directly test how CRC tumoroids influence macrophage polarization, revealing a strong induction of the SPP1+ state in human macrophages derived from circulating monocytes, indicating that carcinoma cells themselves can induce this state in the absence of cancer-associated fibroblasts or a hypoxic environment. Conversely, the ability of SPP1+ macrophages to influence carcinoma cell identity appeared to correlate with expression levels of CD44, the Osteopontin receptor. In tumoroids that were responsive to macrophage co-culture, induction of EMT was observed, consistent with prior reports on pro-metastasis and EMT promoting functions of Osteopontin^34,32^.

Ultimately, this study uncovers shortcomings and advantageous of human patient-derived tumor organoid culture systems, providing a framework for utilizing these systems in colorectal cancer research and therapeutic development.

## Materials and Methods

### Human tissues

Human colorectal cancer specimens were obtained from patients undergoing elective surgery at Hospital of the University of Pennsylvania with written informed consent under the protocol approved by the University of Pennsylvania Institutional Review Board (Protocol number 827759 and PI - Dr. Bryson Katona). For those individuals who consent, after surgical resection, tumors are received in pathology, grossly examined by a pathologist, and a section of tumor and normal adjacent colon is obtained when extra tissue can be provided for research use without compromising patient care. Tissue is then transferred into ice cold phosphate-buffered saline (PBS) and placed on ice until use.

### Tissue dissociation and cell sorting

Tumor and normal colon tissue was dissociated at 37°C using a commercial Human Tissue Dissociation Kit (Miltenyi Order no. 130-095-929) following manufacturer’s instruction. Briefly, 50-100 mg tissue was minced into small pieces using a surgical blade and then incubated with 5 ml dissociation solution for 30 min on a tube rotator at 12 rpm. Tissue fragments settling down at the bottom of the tube are collected and minced again. DNase I (NEB Order no. M0303L) is then to the dissociation solution to the final concentration of 10 U/ml (200X) and incubated at 37°C for another 30 min. The cell pellet is resuspended in 1ml Red Blood Cell Lysis Solution (Sigma Order no. 11814389001) for 4 min at RT and diluted to 15ml using HBSS buffer containing 1% BSA. Any residual fragments are removed by filtering samples just prior to cell sorting through a 35-µm nylon mesh. All centrifugation steps are set to 250 x g for 5 min at 4°C. DAPI (Thermo Fisher Order no. 62248) was used for living versus dead discrimination. Cells were sorted using a Becton Dickinson (BD) FACS Aria controlled by BD FACS DIVA software. FSC-H, FSC-W, SSC-H, SSC-W in combination serve to exclude doublets. Cells were sorted into an Eppendorf protein low-binding tube containing HBSS with 0.04% BSA.

### Single-cell RNA-seq

Sorted cells were immediately processed using a10x Genomics Chromium controller and the Chromium Single Cell 3’ Reagent Kits V3 protocol. 7,000-16,000 cells were loaded for each sample. Cells were partitioned into gel beads, lysed, and barcoded through reverse transcription. cDNA was purified and amplified using appropriate cycle number following the manufacturer’s protocol. Libraries were constructed using 10x Genomics Library Prep Kit. Library quality was checked using Agilent High Sensitivity DNA Kit and Bioanalyzer 2100. Libraries were quantified using dsDNA High-Sensitivity (HS) Assay Kit (Invitrogen) on Qubit fluorometer and the qPCR-based KAPA quantification kit. Libraries were sequenced on an Illumina Nova-Seq 6000 with 28:8:0:87 paired-end format.

### Organoid culture

Organoid culture was carried out following the method previously described ^38^ from fresh surgically resected human colorectal adenocarcinomas or adjacent normal colon. For normal colonic crypt culture, the surface epithelial lining was first removed by surgical scalpel. The remaining tissue was washed vigorously with cold DPBS containing Pen/Strep and Gentamicin for 10 times before incubating with 2.5 mM EDTA (Invitrogen, AM9260G) for 15 min at 4 °C. Crypts were liberated by gently pushing the tissue through a 25 ml serological pipette against the bottom of the tube with a narrow gap in between. Crypts were suspended in Matrigel and dispensed onto 48-well-plate in a 100 ul drop. Tumor sample dissociation followed the protocol detailed in the cell sorting section above with the following modification: instead of enriching the single cell fraction, tissue clusters isolated in the cell strainer were collected and resuspended in Matrigel (BD, 356231) and seeded in a 100 ul drop. The final seeding density is 100 intact crypts per drop. Medium composition is as follows: Advanced Dulbecco’s Modified Eagle’s Medium/F12 (Thermo Fisher Scientific, 12634-010) was supplemented with penicillin/streptomycin (Thermo Fisher Scientific, 15140-122), 10 mM HEPES (Thermo Fisher Scientific, 15630-080), 2 mM GlutaMAX (Thermo Fisher Scientific, 35050-061), 1 X N2 Supplement (Thermo Fisher Scientific, 17502-048), 1 X B-27 Supplement (Thermo Fisher Scientific, 17504-044) to prepare a basal medium. An expansion medium was made by supplementing the basal medium with 10 nM gastrin I (Sigma-Aldrich, G9145-.1MG), 1 mM N-acetylcysteine (Sigma-Aldrich, A9165-5g), 100 ng/ml recombinant mouse Noggin (PeproTech, 250-38), 50 ng/ml recombinant mouse EGF (Thermo Fisher Scientific, PMG8041), 1 ug/ml recombinant human R-spondin1 (R&D, 4645-RS-025), 500 nM A83-01 (Tocris, 2939),10 uM SB 202190 (Sigma-Aldrich, S7067-5MG). The organoids were passaged biweekly by digesting with TyrpLE (Thermo Fisher Scientific, 12604013) for 5 min at 37 °C and then splitting in 1:3-5 ratio. The medium was supplemented with 10 uM Y-27632 (Selleck, S1049) for the first two days of culture after seeding.

### Monocyte isolation and macrophage differentiation

Normal donor human monocytes were procured from the Human Immunology Core at the University of Pennsylvania. Cells were cultured in RPMI-1640 medium (with 10% FBS, 1% Pen/Strep and 2mM glutamine). Cells were treated with 50ng/ml human M-CSF (PEPROTECH, Cat# 300-25-50ug) for 7 days. Then collect cells. Macrophage cell marker, CD16 (BD biosciences, Cat# 335806), was detected by flow cytometry for verification purpose.

### Organoid-macrophage co-culture

Two days before the co-culture procedure, organoids were dissociated into single cells and seeded within Matrigel drops in 12-well-plates. When setting up the co-culture, organoids were released from the Matrigel by incubating with Corning Cell Recovery Solution for 20 min at 4 °C. Intact organoids were mixed with 10^5^ monocyte-derived macrophages and seeded back to the 12-well-plate. The culture was maintained up to 48h and then harvested for analysis.

### Single cell RNAseq data processing

Sequencing reads for the human and mouse samples were first pre-processed with 10x Genomics Cell Ranger pipeline and aligned to the GRCh38 reference and GRCm38 (mm10) reference genome respectively. An initial filtering was performed on the raw gene-barcode matrix output by the Cell Ranger cellranger count function, removing barcodes that have less than 1000 transcripts (quantified by unique molecular identifier (UMI)) and 500 expressed genes (“expressed” means that there is at least 1 transcript from the gene in the cell). Barcodes that pass this filter were considered as cells and were fed into downstream analysis. For samples multiplexed using the TotalSeq-B protocol, cells were demultiplexed by performing Louvain clustering on the UMAP generated with the hashtag count matrix. Gene-barcode UMI count matrix combined from all datasets was size-factor corrected and log transformed to produce a normalized gene expression matrix.

### Dimension reduction, clustering and cell type annotation

We used the VisCello package to generate a series of PCAs and UMAPs for different cell subsets. The processing pipeline was described previously in Zhu et al^39^. Briefly, we applied an “informative feature (IFF) selection” procedure to select genes that have high gini coefficient which indicates the “inequality” (therefore specificity) of the gene’s expression across clusters. Principal component analysis (PCA) was then performed on the IFF-cell matrix, and the top PCs were used as features for the UMAP algorithm. UMAP was computed using the *umap* function in the uwot R package, with “cosine” distance metric, 30 nearest neighbors, and the rest of the parameters as default. Louvain clustering was run on the k-nearest neighbor graph (k = 20) constructed from cell embeddings on the UMAP. We annotated each of the clusters based on comparing cluster-specific differentially expressed genes with known cell-type marker genes based on previous literature. Top DEGs/cell-type markers include: Epitheilal: *EPCAM*, Fibroblast: *VIM,COL1A1,COL1A2*; Myofibroblast: *VIM, ACTA2*; Endothelial: *CDH5,CLDN5,ESAM*; T cells: *CD3D,CD3E*; B cells: *CD19,CD20*; Plasma cells: Immunoglobulin genes, such as *JCHAIN, IGHA, IGHG, IGHM*; Macrophage: *CD68,LYZ*; DC: *CD1C,BATF3*; Mast cells: *TPSAB1, TPSD1, CPA3*. A small cell population from liver/liver metastasis samples form a separate cluster and were broadly annotated as “liver cells” as we do not have enough resolution to distinguish the cell subtypes.

### Differential expression analysis

Differential expression analysis was carried out using the “sSeq” algorithm, with FDR < 0.05 and log2 fold change > 1 as cutoff for differentially expressed genes (DEGs). For DE between liver metastasis and primary tumor shown in Supplemental Figure 2, Mann-Whitney U test was used instead because of the limited cell number and sequencing depth.

### Pathway/signature enrichment analysis

To compute a per-cell enrichment score for each pathway in the Reactome database or cell-type specific signature shown in Figure 3D, S3E, 4H, S4H, 5E, 5F, S5B, 6C, 6G and Supplemental Figure 2C, we utilized the AUCell package. The AUCell package uses the “Area Under the Curve” (AUC) to calculate the enrichment of a particular gene set among top expressed genes of each cell. Quantification of pathway activity and cell-type signature score enables subsequent statistical test, such as ANOVA and Tukey’s test in Figure 6C, and Student’s *t* test to derive up-regulated Reactome pathways in liver metastasis shown in Supplemental Figure 3E.

Gene ontology (GO) enrichment analyses in Figure 3F and 4F were computed using the ClusterProfiler package with q-value cutoff of 0.05 and ontology type “Biological Process” (BP).

### Analysis of condition-specific carcinoma-TME interactions

Receptor/ligand information was obtained from a curated database^40^. To predict receptor-ligand interactions that are specifically induced by the tumor microenvironment, we took advantage of the normal adjacent epithelial cells we collected and used these as the control to derive receptor/ligands that are specifically up-regulated in the carcinoma cells. For each of the receptors/ligands significantly up-regulated (FDR < 0.05), we looked for its counterpart in the non-epithelial TME cells. TME cell types that differentially express the receptor/ligand counterpart were considered to be communicating with the tumor epithelial cell via the receptor-ligand interaction. The receptor-ligand interactions between tumor epithelial cell and TME cells were visualized as a bipartite graph using the ggraph package.

### Identification of macrophage states and FateID analysis

Monocytes/macrophages were first identified using classic monocyte/macrophage marker genes such as *CD14, CD68* and *LYZ*. Louvain clustering was then performed on the subset of cells, revealing large heterogeneity within the monocyte/macrophage cell subset. To annotate the different cell states within this population, we first performed one-vs-rest differential gene expression analysis and derived lists of differentially expressed genes. We then compared the top differentially expressed genes with known marker genes of macrophage states based on previous literature^15,18^ and adopted the commonly used markers to annotate the cell states. Interestingly, expression patterns of three key state marker genes, *IL1B, SPP1* and C1QC partitioned the cell population into three groups, with relatively small overlaps between each other (Figure 4C). The partition of the cells is also consistent with the Louvain clustering result. To probabilistically assign cells to each of the states, we re-derived state-specific genes using differential expression analysis, and performed FateID analysis with these signature genes and default parameters (Figure 4D, E). Pathway enrichment analysis was performed on the state-specific genes, revealing distinct set of pathways being activated in each of the macrophage states (Figure 4F).

### Consensus molecular subtyping of colorectal cancer

Consensus molecular subtypes were called using the CMScaller R package^41^. For each sample, we aggregated its single-cell gene expression counts to obtain a pseudo-bulk expression vector. We run CMScaller with parameter *RNASeq*=TRUE and the rest parameters as default. CMScaller performs classification using nearest template prediction algorithm with pre-defined cancer-cell intrinsic CMS templates (**Supplemental Table 3**).

### CODEX antibody conjugation

Akoya antibodies were purchased preconjugated to their respective CODEX barcode (**Supplemental Table 4**). All other antibodies were custom conjugated to their respective CODEX barcode (**Supplemental Table 4**) according to Akoya’s CODEX user manual using the CODEX Conjugation Kit (Akoya, 7000009). Briefly, 50 μg of carrier-free antibodies (**Supplemental Table 4**) were concentrated by centrifugation in 50kDa MWCO filters (EMD Millipore, UFC505096) and incubated in the antibody disulfide reduction master mix for 30 minutes. After buffer exchange of the antibodies to conjugation solution by centrifugation, addition of conjugation solution and centrifugation, respective CODEX barcodes resuspended in conjugation solution were added to the concentrated antibody and incubated for 2 hours at room temperature. Conjugated antibodies were purified by 3 buffer exchanges with purification solution. 100 μl of antibody storage buffer was added to the concentrated purified antibodies.

### CODEX staining

CODEX staining was done using the CODEX staining kit (Akoya, Cat 7000008) according to Akoya’s CODEX user manual with modifications to include a photobleaching step and overnight incubation in antibodies at 4°C. FFPE samples were sectioned at 5 μm and mounted onto 22 mm x 22 mm coverslips coated with poly-L-lysine (Sigma-Aldrich, P8920) coated according to Akoya’s CODEX user manual. Sample coverslips were heated on a 55°C hot plate for 25 minutes to bake the tissue. Sample coverslips were deparaffinized in xylene and rehydrated in a graded series of ethanol (2 times 100%, 90%, 70%, 50%, 30% and 2 times ddH2O). Antigen retrieval was performed in 1X Tris-EDTA buffer pH 9.0 (Abcam, ab93684) with a pressure cooker for 20 minutes. After equilibrating to room temperature, sample coverslips were washed 2 times with ddH2O and submerged in a 6-well plate containing 4.5% H2O2 and 20mM NaOH in PBS (bleaching solution) for photobleaching. The 6-well plates were sandwiched between two broad-spectrum LED light sources for 45 minutes at 4°C. Sample coverslips are transferred to a new 6-well plate with fresh bleaching solution and photobleached for another 45 minutes at 4°C. Sample coverslips were washed 3 times in PBS and then 2 times in hydration buffer. Sample coverslips were equilibrated in staining buffer for 30 minutes and incubated in the antibodies (Table 1) diluted in staining buffer plus N Blocker, G Blocker, J Blocker, and S Blocker overnight at 4°C. After antibody incubation, sample coverslips were washed 2 times in staining buffer and fixed for 10 minutes in 1.6% paraformaldehyde (Electron Microscopy Sciences, 15710) in storage buffer. Sample coverslips were washed 3 times in PBS and incubated in ice cold methanol for 5 minutes. After incubation in methanol, sample coverslips were washed 3 times in PBS and incubated in final fixative solution for 20 minutes. The sample coverslips were then washed 3 times in PBS and stored in storage buffer.

### CODEX imaging

CODEX reporters were prepared according to Akoya’s CODEX user manual and added to a 96-well plate. The CODEX instrument was set up for a CODEX run according to Akoya’s CODEX user manual using the CODEX instrument manager software. Details on the order of fluorescent CODEX Barcodes and microscope exposure times, can be found on Table 2. Images were taken with a Nikon Plan Apo λ 20X/0.75 objective on a Keyence BZ-X810 fluorescence microscope. Microscope setup was done according to Akoya’s CODEX user manual with a z plane of 11 and z pitch of 1.2 μm. 2 areas of 1.5 mm x 1.1 mm were imaged for patient 86 and patient 92. 3 areas of 2.0 mm x 1.5 mm were imaged for patient 28.

### CODEX image analysis

The CODEX images were processed with the CODEX® processor software, which performs background subtraction, deconvolution, stitching and segmentation with default settings. The segmented data were imported into R, where a two-component gaussian mixture model was fit to the intensity values of each channel. The component with a lower mean was treated as background noise and only signals from the higher component was retained. The corrected intensity values were then log transformed and imported into VisCello^42^ for clustering, cell type annotation and visualization.

### BioRender

Cartoons in Figure1, 6 were created with BioRender.com.

## Acknowledgments

This work was funded by a Translational Center for Excellence award from the Abramson Cancer Center at the University of Pennsylvania. CJL is supported by R01 CA168654. KT is supported by U01 CA243072 and U2C CA233285. NL is supported by R50 CA221841. ATT is supported by R01 CA196299. AR is supported by R01 CA272903 and the HICC P30CA013696. This work was supported by core facilities funded by the NIDDK P30 Center for Molecular Studies in Digestive and Liver Diseases (P30DK050306) at the University of Pennsylvania.

## Figure Legends

**Supplemental Figure 1.**
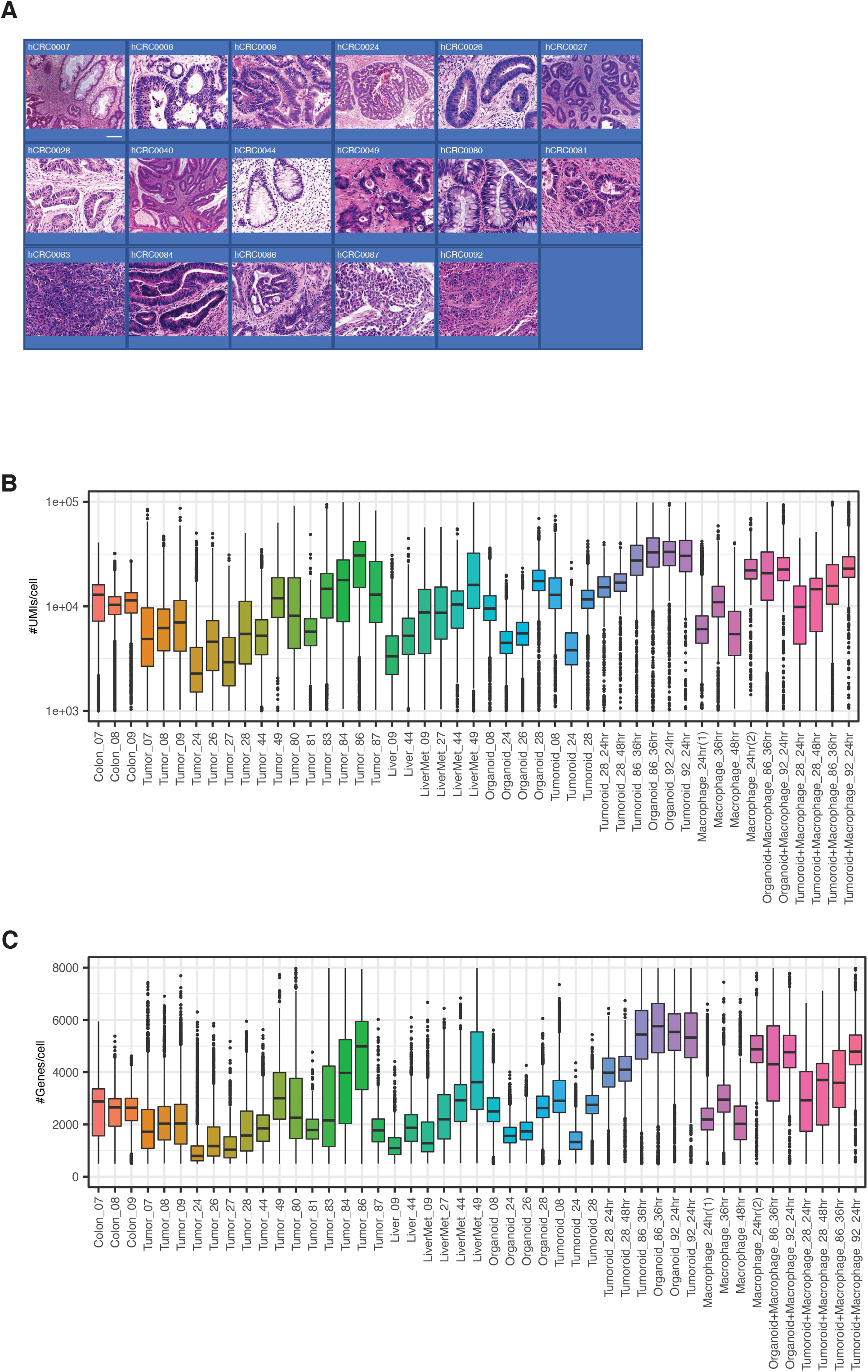
Single cell transcriptomic profiling of primary colorectal cancers and their cultured tumoroid derivatives. (**A**) Hematoxylin/eosin micrographs of primary tumors not included in Figure 1. (**B**) Average number of unique molecular identifiers (UMIs) per cell across scRNA-Seq samples. (**C**) Average number of genes detected per cell across scRNA-Seq samples.

**Supplemental Figure 2.**
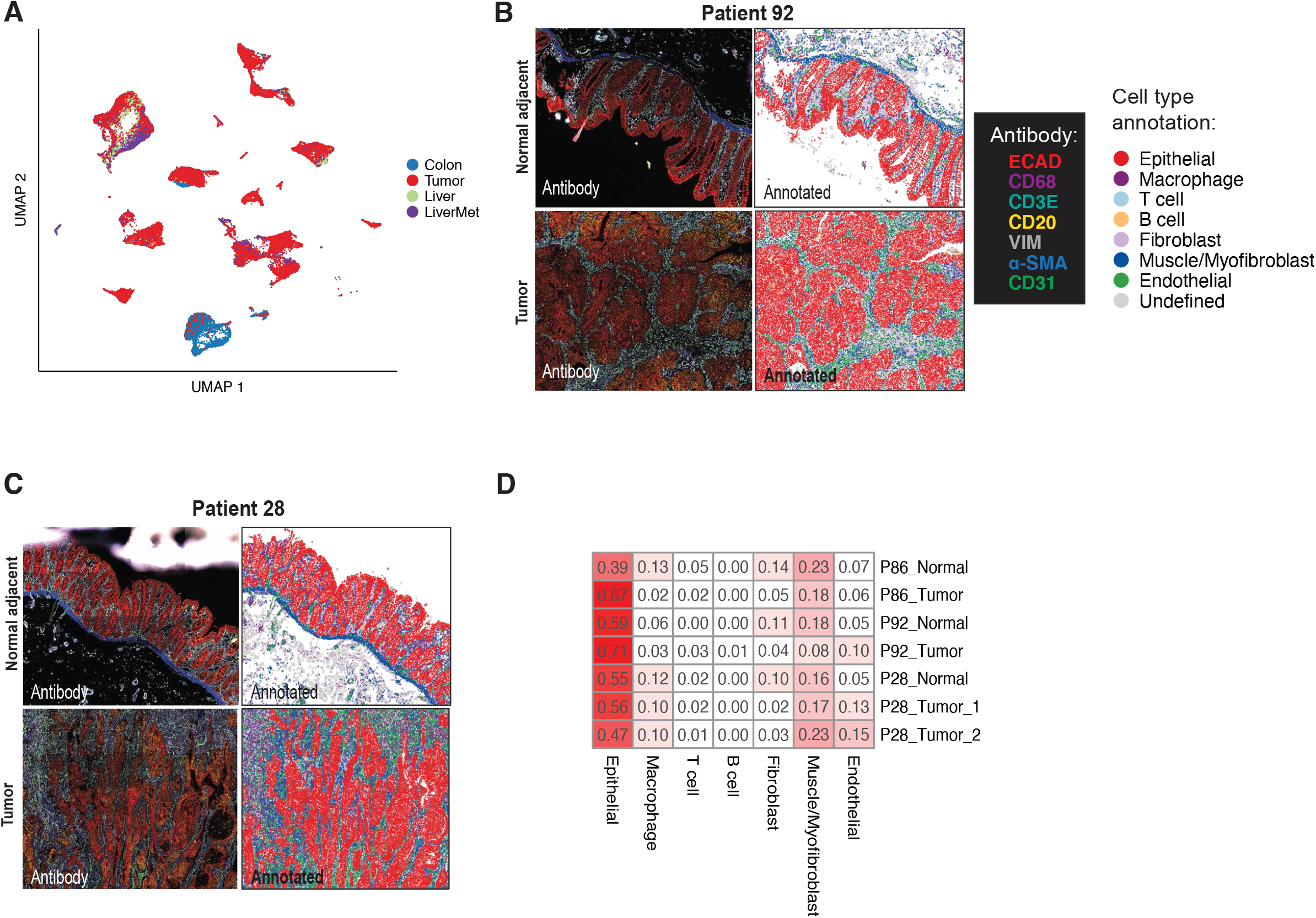
Spatial relationships between carcinoma cells and the TME. (**A**) UMAP as in Figure 2A, highlighting the source type of the cells. (**B, C**) Left panel: CODEX image of normal adjacent tissues and primary tumor from patient 92 (B) and 28 (C), highlighting seven cell-type markers – ECAD, CD68, CD3E, CD20, VIM, α-SMA and CD31. Right panel: post-segmented image colored by cell type. (**D**). CODEX-based cell type composition for samples from patient 86 (Figure 2C), patient 92 (B) and patient 28 (C and for another tumor section not shown here). Value represents proportion of each cell type in the sample normalized by total cell count.

**Supplemental Figure 3.**
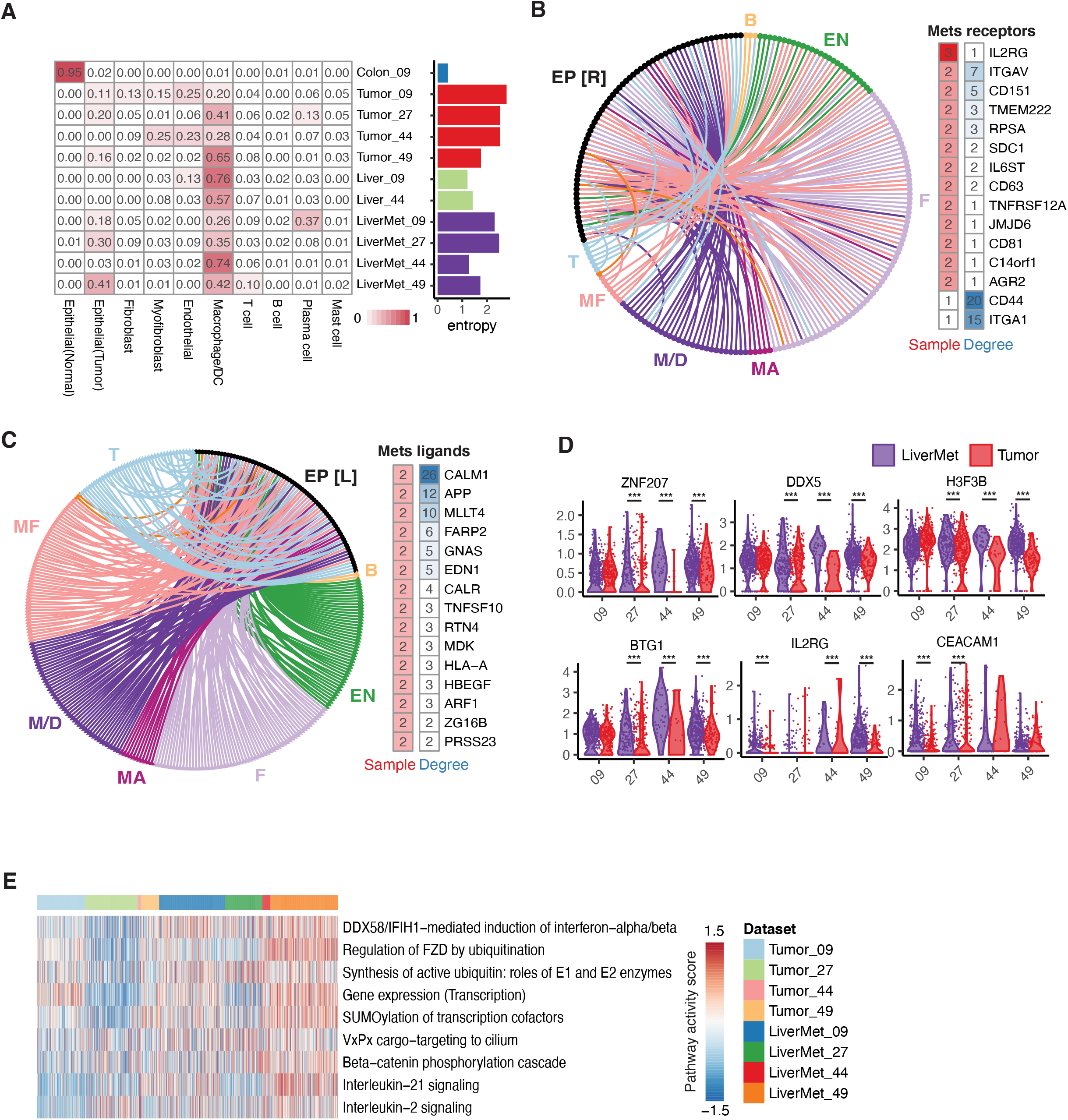
Analysis of liver metastasis of human colorectal cancer. **A**) Cell type composition of each sample. Only patients with liver metastasis data were included. Value represents proportion of each cell type in the sample (normalized by total cell count instead of epithelial cell count as in Figure 2B, due to lack of normal colon epithelial cells in the liver samples). For each sample, entropy was calculated based on the cell type proportion to measure the heterogeneity of TME of each sample. (**B**) Receptor-ligand interaction up-regulated in liver metastasis. Each edge indicates communication between a receptor up-regulated in metastatic tumor epithelial cells compared to primary tumor epithelial cells, and a ligand expressed by microenvironment cells. Receptors were ranked based on the number of patients in which the gene is differentially expressed, and the degree on the communication graph. EP [R]: receptors expressed on epithelial cells; B: B cells; T: T cells; EN: endothelial cells; F fibroblasts; MF: Myofibroblasts; M/D: Macrophage/dendritic cells. MA: mast cells. (**C**) Same as (B), but highlighting the ligands up-regulated in metastatic tumor epithelial cells compared to primary tumor epithelial cells, and corresponding receptors expressed by microenvironment cells. (**D**) Violin plots of expression level of selected genes that show consistently increased expression in liver metastasis compared to primary tumor across multiple patients. (**E**) Top up-regulated Reactome pathways in liver metastasis. Pathway scores for each cell were computed using AUCell package followed by t-test to derive differentially activated pathways.

**Supplemental Figure 4.**
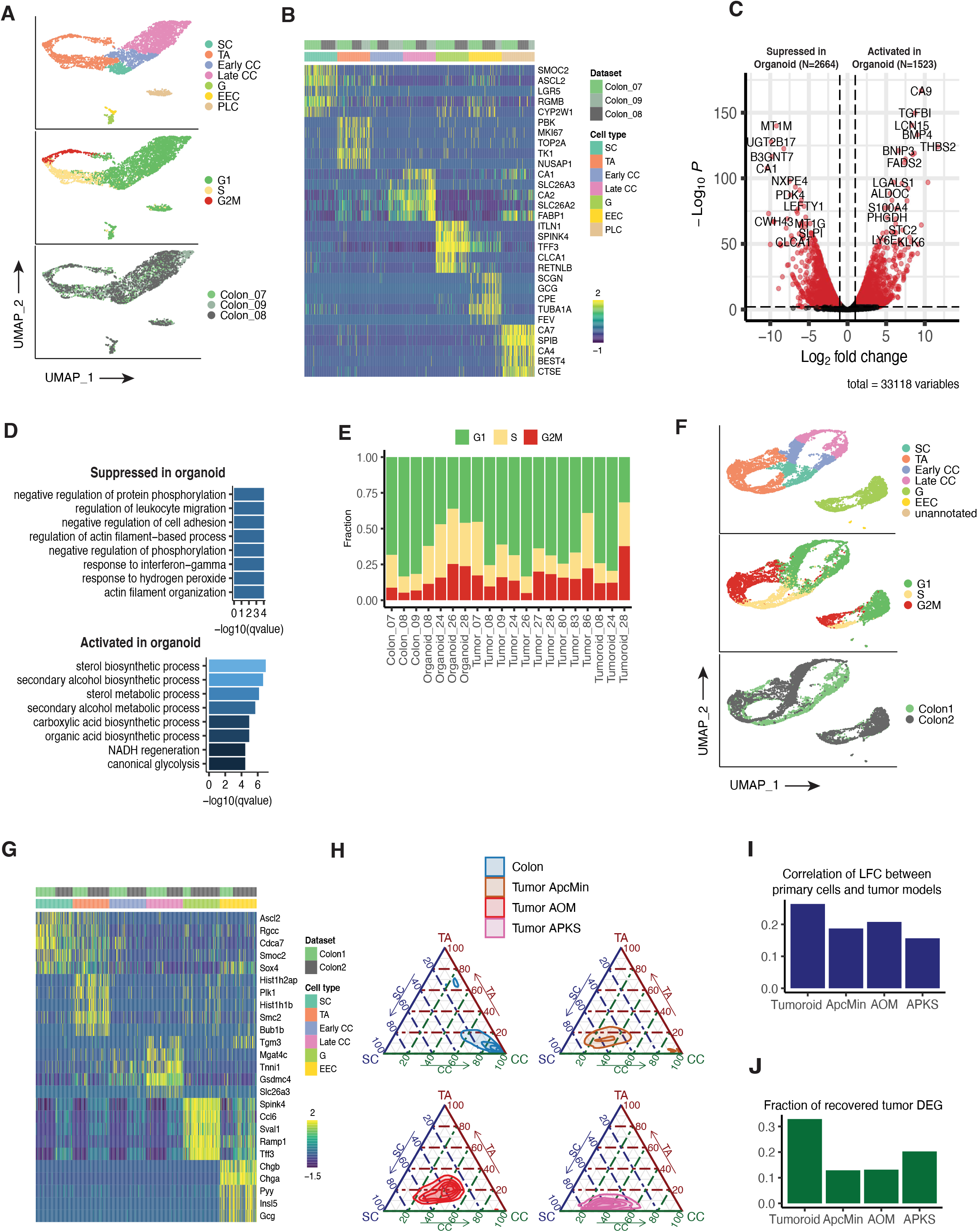
Analysis of human carcinoma and normal epithelial cells *in vivo* and *in vitro* with comparison to mouse models of colorectal cancer. (**A**) UMAP of normal adjacent colon epithelial cells, colored by cell type, cell cycle phase and patient id. SC: stem cells; TA: transit amplifying (TA) cells; Early CC: early colonocytes; Late CC: late colonocytes; G: goblet cells; EEC: enteroendocrine cells; PLC: Paneth-like cells. (**B**) Expression heatmap of top differentially expressed genes of each cell type in human. (**C**) Volcano plot showing genes activated and suppressed in organoid compared to normal adjacent colon tissue based on differential expression analysis. Red point represents genes significantly up- or down-regulated in organoid compared to normal adjacent colon, with FDR ≤ 0.01 and log2 fold change ≥ 1. (**D**) Gene ontology (GO) functional analysis of DEGs activated and suppressed in organoids-log_10_(q-value) of the top significantly enriched GO terms in biological process were plotted. (**E**) Bar plot showing cell cycle phase composition for each dataset. (**F**) UMAP of mouse normal colon, colored by cell type, cell cycle phase, and replicate id. Cells were merged from two separate scRNA-Seq experiments and display some batch effect. However, both batches showed consistent expression of cell-type marker genes. SC: stem cells; TA: transit amplifying (TA) cells; Early CC: early colonocytes; Late CC: late colonocytes; G: goblet cells; EEC: enteroendocrine cells; PLC: Paneth-like cells. (**G**) Expression heatmap of top differentially expressed genes of each cell type in mouse. (**H**) Ternary plot showing distribution of cells in the cell-type-signature space across normal mouse colon and mouse models of colorectal cancer (ApcMin/+, AOM-DSS, and APKS tumoroid implantation models). Cell type signature genes were derived for each epithelial cell types including stem cell (SC), transit amplifying cell (TA) and colonocytes (CC) using differential expression analysis on normal epithelial cells. Signature scores were calculated using the AUCell package, and linearly rescaled to 0 to 100. 2D kernel density estimation was performed to visualize the cell distribution. (**I**) Correlation of LFC between primary human carcinoma cells, human tumoroids and mouse tumor models. (**J**) Bar plot showing fraction of primary-tumor-specific genes that can be recovered using various mouse tumor models.

**Supplemental Figure 5.**
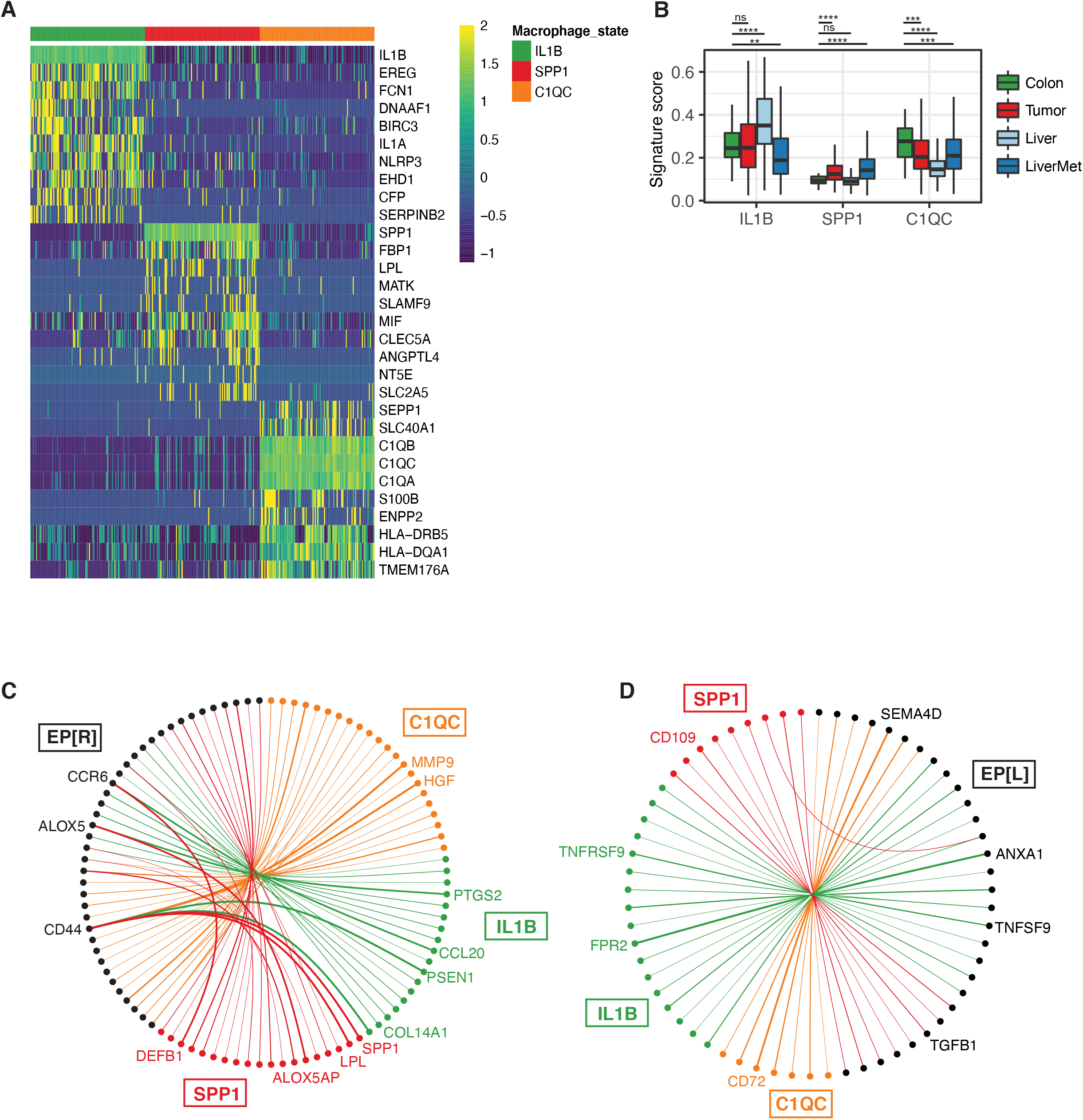
Signature genes of macrophage states. (**A**) Gene expression heatmap of top differentially expressed genes from macrophages in the IL1B, SPP1 and C1QC states. Expression values were log-normalized and z-scored. (**B**) Distribution of signature score of macrophage states across different sample types. Two-sided student’s t test was performed between colon and tumor, colon and liver and colon and liver metastasis. **p≤0.01; ***p≤0.001; ****p≤0.0001; ns, p>0.05. **C**) Receptor-ligand interactions between primary tumor and macrophage subpopulations. Each edge indicates a predicted interaction between a receptor up-regulated in primary tumor carcinoma cells compared to normal adjacent epithelial cells, and a ligand differentially expressed between the macrophage states. Edge widths indicate the number of patients (samples) in which the receptor is significantly up-regulated. EP [R]: receptors expressed on epithelial cells. (**D**) Same as C, but highlighting the ligands up-regulated in primary tumor carcinoma cells compared to normal adjacent epithelial cells, and corresponding receptors differentially expressed by macrophage states. EP [L]: ligands expressed on epithelial cells.

## References

1. Rawla, P., Sunkara, T. & Barsouk, A. Epidemiology of colorectal cancer: incidence, mortality, survival, and risk factors. Prz Gastroenterol 14, 89–103 (2019).

2. Stoffel, E. M. & Murphy, C. C. Epidemiology and Mechanisms of the Increasing Incidence of Colon and Rectal Cancers in Young Adults. Gastroenterology 158, 341–353 (2020).

3. Sato, T. et al. Single Lgr5 stem cells build crypt-villus structures in vitro without a mesenchymal niche. Nature 459, 262–265 (2009).

4. Sato, T. et al. Long-term expansion of epithelial organoids from human colon, adenoma, adenocarcinoma, and Barrett’s epithelium. Gastroenterology 141, 1762–1772 (2011).

5. Ganesh, K. et al. A rectal cancer organoid platform to study individual responses to chemoradiation. Nat Med 25, 1607–1614 (2019).

6. Narasimhan, V. et al. Medium-throughput Drug Screening of Patient-derived Organoids from Colorectal Peritoneal Metastases to Direct Personalized Therapy. Clin Cancer Res 26, 3662–3670 (2020).

7. Ooft, S. N. et al. Patient-derived organoids can predict response to chemotherapy in metastatic colorectal cancer patients. Sci Transl Med 11, (2019).

8. Bruun, J. et al. Patient-Derived Organoids from Multiple Colorectal Cancer Liver Metastases Reveal Moderate Intra-patient Pharmacotranscriptomic Heterogeneity. Clin Cancer Res 26, 4107–4119 (2020).

9. Vlachogiannis, G. et al. Patient-derived organoids model treatment response of metastatic gastrointestinal cancers. Science 359, 920–926 (2018).

10. Yao, Y. et al. Patient-Derived Organoids Predict Chemoradiation Responses of Locally Advanced Rectal Cancer. Cell Stem Cell 26, 17-26.e6 (2020).

11. Neal, J. T. et al. Organoid Modeling of the Tumor Immune Microenvironment. Cell 175, 1972-1988.e16 (2018).

12. Rogoz, A., Reis, B. S., Karssemeijer, R. A. & Mucida, D. A 3-D enteroid-based model to study T-cell and epithelial cell interaction. Journal of Immunological Methods 421, 89–95 (2015).

13. Noel, G. et al. A primary human macrophage-enteroid co-culture model to investigate mucosal gut physiology and host-pathogen interactions. Sci Rep 7, 45270 (2017).

14. Ye, W., Luo, C., Li, C., Huang, J. & Liu, F. Organoids to study immune functions, immunological diseases and immunotherapy. Cancer Letters 477, 31–40 (2020).

15. Zhang, L. et al. Single-Cell Analyses Inform Mechanisms of Myeloid-Targeted Therapies in Colon Cancer. Cell 181, 442-459.e29 (2020).

16. Liu, Y. et al. Immune phenotypic linkage between colorectal cancer and liver metastasis. Cancer Cell 40, 424-437.e5 (2022).

17. Chen, B. et al. Differential pre-malignant programs and microenvironment chart distinct paths to malignancy in human colorectal polyps. Cell 184, 6262-6280.e26 (2021).

18. Li, X., Zhang, Q., Chen, G. & Luo, D. Multi-Omics Analysis Showed the Clinical Value of Gene Signatures of C1QC+ and SPP1+ TAMs in Cervical Cancer. Front. Immunol. 12, 694801 (2021).

19. Qi, J. et al. Single-cell and spatial analysis reveal interaction of FAP+ fibroblasts and SPP1+ macrophages in colorectal cancer. Nat Commun 13, 1742 (2022).

20. Klement, J. D. et al. An osteopontin/CD44 immune checkpoint controls CD8+ T cell activation and tumor immune evasion. Journal of Clinical Investigation 128, 5549–5560 (2018).

21. Guinney, J. et al. The consensus molecular subtypes of colorectal cancer. Nat Med 21, 1350–1356 (2015).

22. Lau, W. de. WNT signaling in the normal intestine and colorectal cancer. Front Biosci 12, 471 (2007).

23. de Sousa e Melo, F. et al. A distinct role for Lgr5+ stem cells in primary and metastatic colon cancer. Nature 543, 676–680 (2017).

24. DeNardo, D. G. & Ruffell, B. Macrophages as regulators of tumour immunity and immunotherapy. Nat Rev Immunol 19, 369–382 (2019).

25. Spranger, S., Dai, D., Horton, B. & Gajewski, T. F. Tumor-Residing Batf3 Dendritic Cells Are Required for Effector T Cell Trafficking and Adoptive T Cell Therapy. Cancer Cell 31, 711-723.e4 (2017).

26. Nizzoli, G. et al. Human CD1c+ dendritic cells secrete high levels of IL-12 and potently prime cytotoxic T-cell responses. Blood 122, 932–942 (2013).

27. Zhang, Q. et al. Landscape and Dynamics of Single Immune Cells in Hepatocellular Carcinoma. Cell 179, 829-845.e20 (2019).

28. Szpor, J. et al. Dendritic Cells Are Associated with Prognosis and Survival in Breast Cancer. Diagnostics 11, 702 (2021).

29. Swiecki, M. & Colonna, M. The multifaceted biology of plasmacytoid dendritic cells. Nat Rev Immunol 15, 471–485 (2015).

30. Wei, J. et al. Characterizing Intercellular Communication of Pan-Cancer Reveals SPP1+ Tumor-Associated Macrophage Expanded in Hypoxia and Promoting Cancer Malignancy Through Single-Cell RNA-Seq Data. Front. Cell Dev. Biol. 9, 749210 (2021).

31. Pitarresi, J. R. et al. PTHrP Drives Pancreatic Cancer Growth and Metastasis and Reveals a New Therapeutic Vulnerability. Cancer Discovery 11, 1774–1791 (2021).

32. Zhao, H. et al. The role of osteopontin in the progression of solid organ tumour. Cell Death Dis 9, 356 (2018).

33. Goltsev, Y. et al. Deep Profiling of Mouse Splenic Architecture with CODEX Multiplexed Imaging. Cell 174, 968-981.e15 (2018).

34. Amilca-Seba, K., Sabbah, M., Larsen, A. K. & Denis, J. A. Osteopontin as a Regulator of Colorectal Cancer Progression and Its Clinical Applications. Cancers 13, 3793 (2021).

35. Tuveson, D. & Clevers, H. Cancer modeling meets human organoid technology. Science 364, 952–955 (2019).

36. Dijkstra, K. K. et al. Generation of Tumor-Reactive T Cells by Co-culture of Peripheral Blood Lymphocytes and Tumor Organoids. Cell 174, 1586-1598.e12 (2018).

37. Zhang, L. et al. Single-Cell Analyses Inform Mechanisms of Myeloid-Targeted Therapies in Colon Cancer. Cell 181, 442-459.e29 (2020).

38. Fujii, M., Matano, M., Nanki, K. & Sato, T. Efficient genetic engineering of human intestinal organoids using electroporation. Nat Protoc 10, 1474–1485 (2015).

39. Zhu, Q. et al. Developmental trajectory of prehematopoietic stem cell formation from endothelium. Blood 136, 845–856 (2020).

40. Shao, X. et al. CellTalkDB: a manually curated database of ligand–receptor interactions in humans and mice. Briefings in Bioinformatics 22, bbaa269 (2021).

41. Eide, P. W., Bruun, J., Lothe, R. A. & Sveen, A. CMScaller: an R package for consensus molecular subtyping of colorectal cancer pre-clinical models. Sci Rep 7, 16618 (2017).

42. Packer, J. S. et al. A lineage-resolved molecular atlas of C. elegans embryogenesis at single-cell resolution. Science 365, eaax1971 (2019).

43. Paternoster, R., Brame, R., Mazerolle, P. & Piquero, A. USING THE CORRECT STATISTICAL TEST FOR THE EQUALITY OF REGRESSION COEFFICIENTS. Criminology 36, 859–866 (1998).

44. Aibar, S. et al. SCENIC: single-cell regulatory network inference and clustering. Nat Methods 14, 1083–1086 (2017).

